# Stratification of human gut microbiomes by succinotype is associated with inflammatory bowel disease status

**DOI:** 10.1101/2023.11.21.568118

**Authors:** Laura Anthamatten, Philipp Rogalla von Bieberstein, Carmen Menzi, Janina N. Zünd, Christophe Lacroix, Tomas de Wouters, Gabriel E. Leventhal

## Abstract

The human gut microbiome produces and consumes a variety of compounds that interact with the host and impact health. Succinate is of particular interest as it intersects with both host and microbiome metabolism. However, which gut bacteria are most responsible for the consumption of intestinal succinate is poorly understood. Here, we build upon an enrichment-based whole fecal sample culturing approach and identify two main bacterial taxa that are responsible for succinate consumption in the human intestinal microbiome, *Phascolarctobacterium* and *Dialister*. These two taxa have the hallmark of a functional guild and are strongly mutual exclusive across over 20,000 fecal samples in nearly 100 cohorts and can thus be used to assign a robust ‘succinotype’ to an individual. We show that they differ with respect to their rate of succinate consumption *in vitro* and that this is associated with higher concentrations of fecal succinate. Finally, individuals suffering from inflammatory bowel disease (IBD) are more likely to have the *Dialister* succinotype compared to healthy subjects. The functionally meaningful classification of human intestinal microbiota based on ‘succinotype’ thus builds a bridge between microbiome function and IBD pathophysiology related to succinate.

## Introduction

The intestinal microbiome interacts with its host is through the production and consumption of physiologically relevant metabolites^1^. This overall microbiome metabolic activity emerges from the individual activity of the member microbes, and thus the overall metabolic output can vary depending on the specific microbiome composition^2^. Screening for and controlling the specific microbes that are the drivers of these activities is a promising diagnostic and intervention target. However, which specific microbe in a microbiome is the key responsible for a specific function remains largely unknown.

Some metabolites—like succinate—are produced and consumed by both the microbiome and the host. On the host side, succinate is a key intermediate of the tricarboxylic acid cycle and thus intricately related to host metabolic homeostasis. On the microbiome side, succinate is an intermediate product of anaerobic carbohydrate fermentation and thus related to microbial energy production^3^. This intersection of host and microbiome metabolism poses a challenge for host regulation: while the host might attempt to regulate how succinate is produced and used, its regulatory control does not expand to the microbiome. As a result, disruptions in how succinate is produced and consumed by the microbiome can have a multifaceted impact on the host. In a healthy gut, succinate is rapidly converted into propionate^4–6^, which in turn is readily absorbed by the host epithelium^7^. However, elevated concentrations of succinate measured in human feces have been associated with intestinal inflammation^8–11^, suggesting a pro-inflammatory effect of excess circulating succinate. Consequently, microbes that metabolize succinate have been suggested to alleviate this inflammatory effect^12,13^.

Only few bacteria are known to anaerobically consume succinate. Various intestinal bacteria—including for example many *Bacteroidaceae*—produce propionate from sugars via the succinate pathway^14,15^, and in some cases export the intermediate succinate due to the low energetic yield of converting succinate to propionate^16^. But these succinate producers have generally not been observed to take up and convert succinate when supplied extracellularly. Succinate can also be otherwise utilized by select bacteria. For example, *Veillonella parvula* can decarboxylate succinate during lactate consumption to produce propionate and increase growth yield^17^, and *Clostridioides difficile* can convert succinate to butyrate to rebalance NADH. However, a substantial consumption of extracellular succinate has only been demonstrated for isolates of the *Negativicutes*, including *Phascolarctobacterium* spp.^4,18^ and *Dialister* spp.^19–21^. Overall, the degree to which these different taxa and pathways are active in the human intestine remains poorly understood.

Here, we aimed to identify and characterize the key bacteria involved in human intestinal succinate consumption. We previously showed that some human fecal microbiomes were able to consume succinate within 48 hours of *in vitro* cultivation, and that this mapped to the presence or absence of certain *Negativicutes* bacteria^22^. We first expanded upon this approach to differentiate between succinate consumption that takes more than 48 h and an overall absence of the function. We observed that all fecal microbiomes had the capacity to consume succinate, but did differ in the rate at which they did so. We then verified that the succinate consumption rate in pure culture mapped to the consumption rate of the whole fecal sample. Our data suggest that bacteria from the genera *Phascolarctobacterium* and *Dialister* are the dominant succinate consumers in the human GI tract, but that *Phascolarctobacterium* converts succinate to propionate significantly more rapidly than *Dialister*. We then analyzed publicly available cohorts of human fecal microbiota to show that *Phascolarctobacterium* and *Dialister* are typically mutually exclusive in human microbiomes, and that IBD patients significantly more likely to have *Dialister* as their dominant succinate consumer compared to healthy individuals. We thus propose that the slower rate of succinate consumption by *Dialister* in the human intestine could be an important contributor to the pathogenesis of intestinal inflammation.

## Results

### Human gut microbiota differ in their ability and rate to metabolize succinate in vitro

To understand succinate consumption in complex intestinal microbiota, we performed *in vitro* enrichments of feces from 13 different human donors akin to what is described in Anthamatten *et al*. [22]. Briefly, we inoculated diluted fecal samples in triplicate into a defined base medium either supplemented with 30 mM of succinate as main carbon source or a non-supplemented control, and measured (i) how much of succinate was consumed after 2 and 7 days of strict anaerobic cultivation and (ii) what metabolites were produced in return (Figure 1a). Because of the stochasticity with respect to diluted inoculum composition, we analyzed each replicate independently.

**Figure 1:**
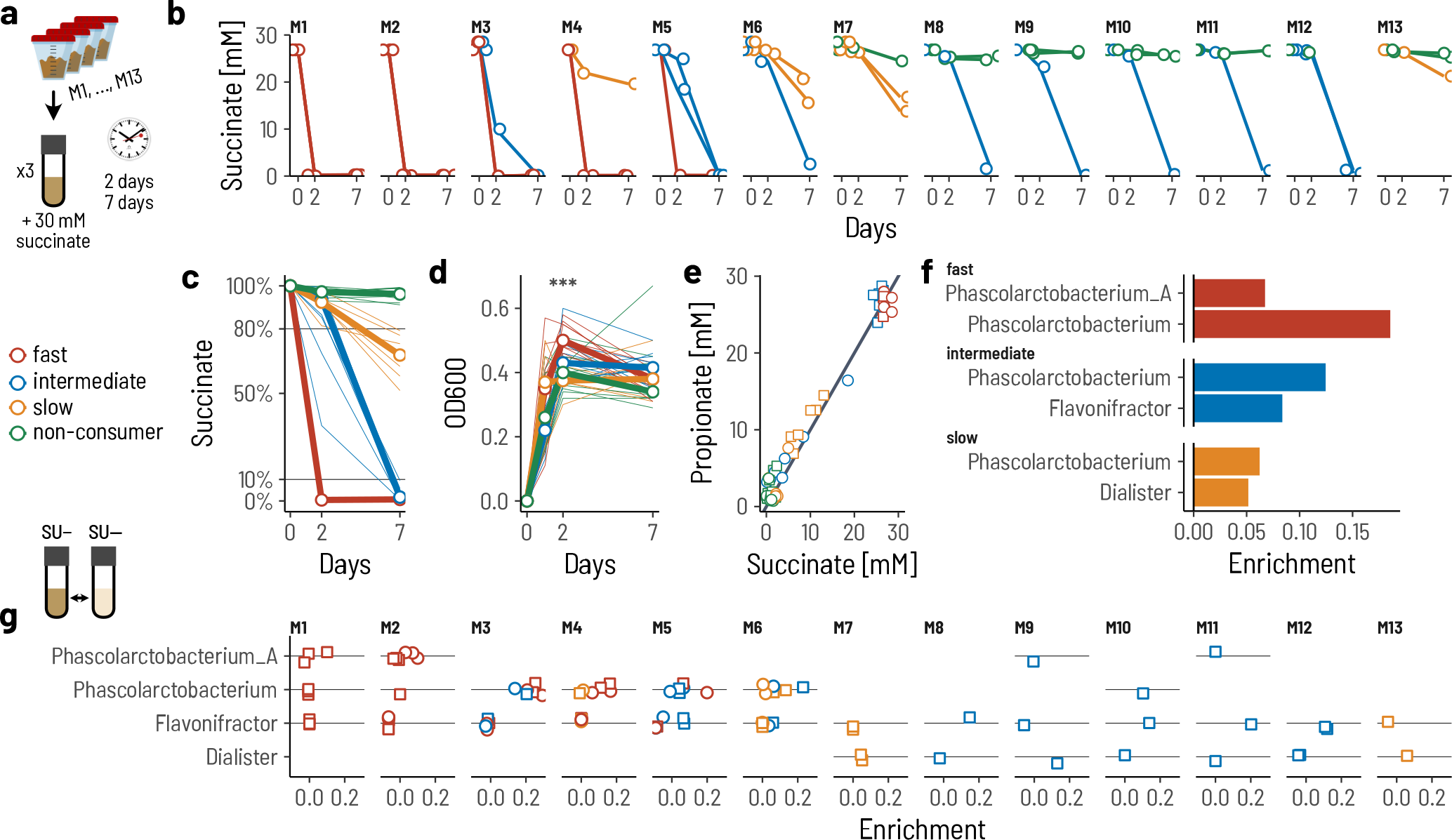
Fecal microbiomes consume succinate at different rates. **a**. We performed enrichment cultures of whole fecal microbiota from 13 different donors in a basal medium supplemented with 30 mM succinate as the primary carbon source. The triplicate cultures were sampled after 2 and 7 days, respectively. **b**. The supplied succinate is consumed differently across fecal microbiota and replicates. **c**. We classify a culture as fast (red), intermediate (blue), or slow (orange) consumer, or non-consumer (green) of succinate. **d**. The increase in optical density (OD) at day 2 was highest for the fast consuming cultures. **e**. The consumed succinate was converted to propionate at a molar ratio of 1:1. **f**. The enrichment of three genera were associated with succinate consumption. **g**. Each fecal microbiome had a distinct signature of putative succinate consuming bacteria.

The time required to consume the supplied succinate differed between enrichment cultures (Figure 1b), with all of the supplied succinate consumed within 48 h in some enrichments and no succinate consumed in others after 7 days. We thus classified the enrichments into four categories as a function of their succinate consumption: ‘fast’, ‘intermediate’, ‘slow’, and ‘non-consumers’ (Figure 1c). Fast enrichments consumed >90% of the supplied succinate within the first 48 h of cultivation (n = 11/39), intermediate enrichments consumed >90% within 7 d (10/39), slow enrichments consumed between 20% and 90% within 7 d (6/39), and non-consuming enrichments consumed <20% within 7 d (12/39). Enrichments inoculated with the same fecal sample were generally consistent in terms of category, with M1 to M4 fast, M5 intermediate, M6 and M7 slow, and M8 to M13 non-consumers. This suggests that specific properties of the fecal sample determine the category of succinate consumption, for example specific taxa or different metabolic pathways. The optical density at 48 h was higher in the fast enrichments compared to the other categories (Figure 1d; Kruskal-Wallis test, *p* = 0.0004), further indicating that different bacteria or even pathways might be associated with the category of succinate consumption. However, succinate was converted to propionate at a molar ratio of 1:1 across all enrichments (Figure 1e) as expected from the succinate pathway^14^, suggesting that this same pathway was ‘in use’ across fecal microbiomes.

We thus next set out to determine which bacterial taxa were performing the conversion of succinate to propionate across enrichments. To this end we performed 16S amplicon sequencing of the succinate enrichment cultures (SU+) and the control cultures (SU-), and computed the differential increase of each genus in SU+ compared to the SU-(see Methods).

Four bacterial genera were significantly associated with the different consumption categories. We performed a linear regression of enrichment in SU+ versus SU- and identified those genera that were significantly associated with at least one of the categories (Supplementary Figure S1). *Phascolarctobacterium* and *Phascolarctobacterium_A* were most strongly associated with the fast category, *Flavonifractor* with the intermediate category, and *Dialister* with the slow category (Figure 1f). Of the four identified succinate consuming taxa, only one was typically dominant in any one specific enrichment (Figure 1g). These data imply that these four genera are most likely those that are responsible for succinate consumption in the enrichments.

Representative genomes of each of these four genera all contained the gene cluster for succinate to propionate conversion first described in *Veillonella parvula*, starting from the succinate-CoA transferase to the methylmalonyl-CoA decarboxylase (Supplementary Figure S2a). The methylmalonyl-CoA decarboxylase subunit alpha (mmdA) has previously been used as a marker gene for the succinate pathway^14^. We wanted to know whether mmdA gene similarity was a good predictor for succinate consumption. To answer this, we reconstructed the phylogenetic tree of mmdA sequences from GTDB and tested a selection of isolates from along the tree for their ability to consume succinate (Supplementary Figure S2b). All the four genera identified in the enrichments could consume succinate in monoculture in the same *in vitro* conditions (Supplementary Figure S2c). However, none of the other tested isolates consumed meaningful amounts of succinate, despite closely related mmdA genes (Supplementary Figure S2c). This suggests that the presence of a homologous mmdA gene— or even the complete succinate pathway as in many *Bacteroidetes*—is not sufficient to confer the ability to consume extracellular succinate in the tested conditions. Taken together, these results suggest that that extracellular succinate consumption in human fecal samples is constrained to very few taxa that include the four genera identified here.

### Intestinal succinate-consuming bacteria differ in their succinate conversion rate

The *in vitro* enrichments essentially test for the competitive ability of the bacteria that comprise the fecal microbiota for succinate. The outcome of such a competition is influenced by two key factors: (i) the *per capita* rate at which the taxa consume succinate, and (ii) the population size of each taxon in the inoculum.

To test whether the identified taxa differ in their *per capita* succinate consumption rate, we performed *in vitro* cultures of eleven representative isolates in a growth medium supplemented with 80 mM succinate and measured the decrease in succinate concentration and the resulting bacterial growth over time (Figure 2a). The representative isolates included two from the genus *Dialister* (*D. hominis* and *D. invisus*), four from *Phascolarctobacterium* (4x *P. faecium*), one from *Phascolarctobacterium_A* (*P. succinatutens*), and three from the genus *Flavonifractor* (3x *F. plautii*). We then estimated the succinate consumption rates of each isolate by deriving a substrate consumption and growth model and subsequently fitting it to the succinate concentration and optical density data in a Bayesian framework (Figure 2a and Methods).

**Figure 2:**
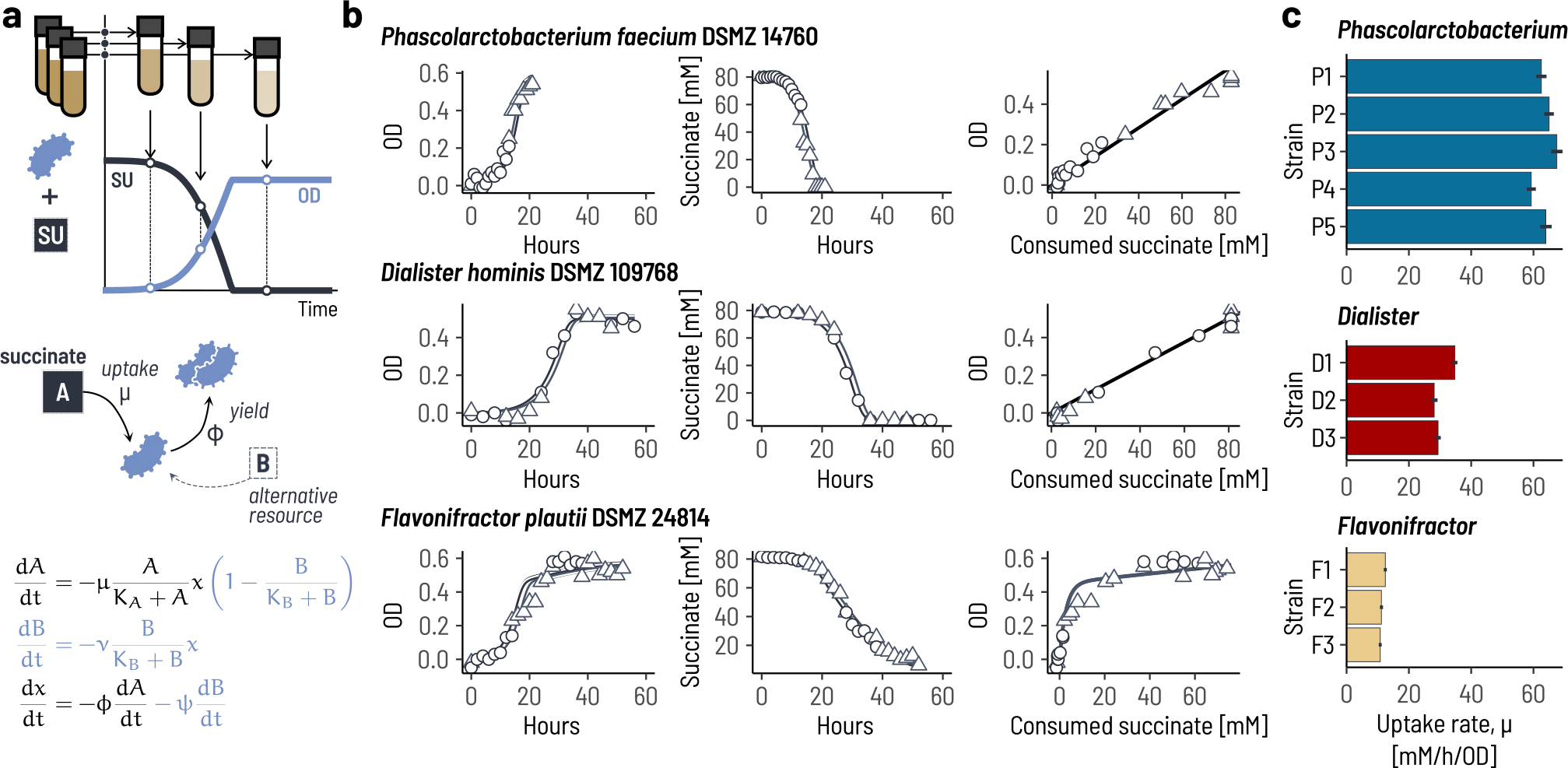
Succinate-consuming gut bacteria differ in their succinate uptake rate. **a**. To estimate the rate at which different representative isolates consume succinate, we performed replicate cultures of a panel of eleven isolates in media supplemented with 80 mM of succinate as the primary carbon source. Each replicate culture was destructively sampled at a different time point between 0 and 60 hours. We then estimated the succinate uptake rate of by fitting a mathematical model of succinate uptake and growth to the data. We accounted for the observed diauxic growth of *Flavonifractor* sp. by adding a second (unobserved) resource to the model (blue terms). **b**. Experimental data and model fits for one representative isolate of each genus. For each isolate, two biological replicate ‘sets’ were inoculated (circles and triangles). Initial bacterial concentrations, x_0_, and succinate concentrations, A_0_, were estimated separately for each replicate set and are shown as separate lines. Data for *Flavonifractor* isolates used the diauxic model. **c**. Posterior mean estimates (bars) and 90% highest-probability density intervals (black lines) for the eleven strains. P1: *P. faecium* DSMZ 14760; P2: *P. faecium* PB-SDVAP; P3: *P. faecium* PB-SJWFW; P4: *P. faecium* PB-SPUPY; P5: *P. succinatutens* DSMZ 22533; D1: *D. hominis* DSMZ 109768; D2: *D. invisus* PB-SARUR; D2: *D. succinatiphilus* DSMZ 21274; F1: *F. plautii* DSMZ 24814; F2: *F. plautii* PB-SCBYV; F3: *F. plautii* PB-SSJQB.

The mathematical model provided a good fit to the experimental data for the *Dialister, Phascolarctobacterium*, and *Phascolarctobacterium_A* isolates (Figure 2b and Supplementary Figure S3). This confirms that these bacteria directly use the energy from converting succinate to propionate for growth. In contrast, the model was not a good fit to the data for *Flavonifractor* for which we observed diauxic growth with a first phase without appreciable succinate consumption (Supplementary Figure S4). To account for this diauxie, we expanded the model to include a second preferred but unobserved growth substrate (Figure 2a). Only once this substrate was depleted does succinate consumption start. This updated model proved a much better fit to the data (Figure 2b and Supplementary Figure S4). With the estimated model parameters at hand, we then compared the strains based on their *per capita* succinate consumption rate.

The estimated succinate uptake rates were consistent within genera but differed strongly between genera (Figure 2c). *Phascolarctobacterium* strains consumed succinate at twice the rate compared to *Dialister* strains, with 63.7 mM*/*h*/*OD on average and 30.7 mM*/*h*/*OD on average, respectively. This translates to longer times required to consume all of the supplemented succinate for the *Dialister* cultures compared to the *Phascolarctobacterium* cultures given equal inoculum densities. *Flavonifractor* strains had an even lower uptake rate, with 11.5 mM*/*h*/*OD on average. However, this did not translate to substantially longer times required to consume the supplied succinate because the *Flavonifractor* cultures first grew on an alternative preferred resource and thus initiate succinate consumption at substantially larger cell densities. This can explain why *Flavonifractor* is more strongly enriched in the fecal microbiomes M8-M12 compared to *Dialister* despite slower *per capita* uptake rate. Overall, these results confirm that the observed differences in the rate of succinate consumption between the whole fecal microbiomes can be mapped to differences in uptake rates of the succinate consuming bacteria.

Having demonstrated that the succinate consumption rate differs between taxa, we next asked to what degree the starting abundances of the succinate consumers might have impacted the overall consumption rate. We thus quantified the relative abundance of the four genera in the thirteen fecal microbiota.

The three genera *Phascolarctobacterium, Phascolarctobacterium_A*, and *Dialister* followed a different compositional pattern than *Flavonifractor* (Figure 3a). The former three all had a bimodal abundance distribution in the fecal samples and were either present at 1–7% or otherwise undetectable. In contrast, *Flavonifractor* was detected in all thirteen fecal microbiota at a consistent abundance of 0.01–0.4%. Because *Flavonifractor* only consumes succinate as a secondary preference, we hypothesized that the bimodal prevalence of the three ‘primary’ consuming genera was the result of mutual exclusion from strong substrate competition. If this was the case, then each fecal sample should only harbor one of the three consumers. Indeed, each of the thirteen fecal microbiota only had one dominant primary succinate consumer (Figure 3b), in most cases with full mutual exclusion (at the sensitivity of sequencing), and this pattern of mutual exclusion also occurred at the species (Figure 3c) and ASV level (Supplementary Figure S5).

**Figure 3:**
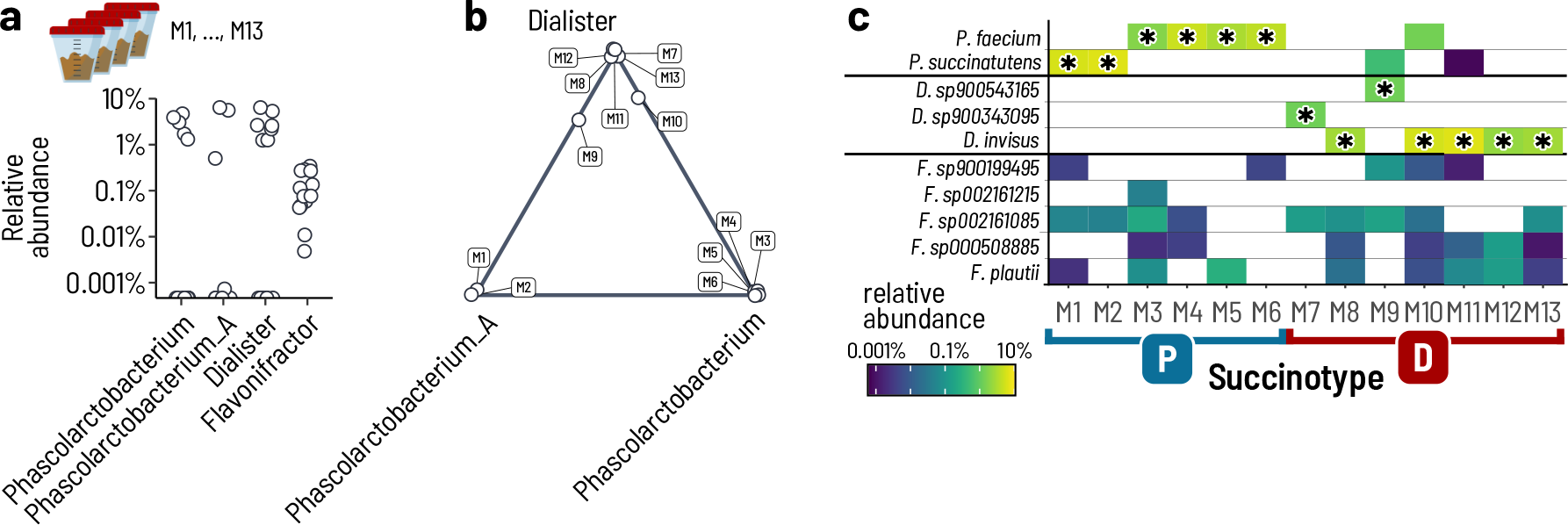
Human fecal microbiota can be classified into ‘succinotypes’ based on their dominant succinate consuming bacterium. **a**. Relative abundances of the four identified succinate consuming genera in the fecal microbiota. **b**. The three genera *Phascolarctobacterium,Phascolarctobacterium_A*, and *Dialister* are strongly mutually exclusive. The position in the ternary diagram shows to the relative abundance of the three genera in each of the fecal microbiota. Points in the corners indicate full mutual exclusivity. **c**. Relative abundances of the succinate consuming taxa in the fecal microbiota resolved at the species level. The asterisk indicates the dominant species in a fecal microbiota. We assign fecal microbiota with a dominant *Phascolarctobacterium,Phascolarctobacterium_A* to a succinotype ‘P’ and those with a dominant *Dialister* to a succinotype ‘D’.

The clear association between the dominant primary succinate consumer and inoculum microbiota indicates that different human intestinal microbiota might be well-classified by the identity of their primary succinate degrader: their ‘succinotype’. Based on the patterns observed here, we introduce the ‘P’ and ‘D’ succinotypes for fecal microbiota that have either *Phascolarctobacterium*/*Phascolarctobacterium_A* or *Dialister*, respectively (Figure 3c). For the P succinotype, we grouped the two genera together because of both their phylogenetic proximity and similarity in *per capita* uptake rate. We next asked whether such a classification of human microbiomes into succinotypes was robust beyond the 13 tested fecal microbiota.

### Succinotypes appear broadly across human cohorts and are associated with disease

To test whether our classification of succinotypes was generalizable to fecal human microbiota more broadly, we analyzed nine publicly available cohorts of 16S amplicon human fecal microbiome data comprising a total of 11,885 samples^23–27^. Furthermore, we also analyzed 85 cohorts with shotgun metagenomic data from the curatedMetagenomicData collection^28^. We used a consistent taxonomic classification based on the Genome Taxonomy Database (GTDB r95)^29^.

The majority of individuals across all cohorts had a well-defined succinotype. Most samples in the amplicon data (91.3%) had detectable abundances of either *Dialister* or *Phascolarctobacterium*, with slightly lower proportions in the shotgun data (76.9%). In order to robustly assign a succinotype, we set either a threshold of at least 10 reads assigned to either D or P (when counts were available) or otherwise a combined relative abundance of at least 0.01%. We retained 8,911 samples after pruning multiple samples from the same subject. We then computed the abundance fraction, *r* = *x*_*D*_*/*(*x*_*D*_ +*x*_*P*_), where the *x* are the abundances of D and P, respectively. The vast majority of samples had *r* values that were close to 0 or 1, respectively (Figure 4a, Supplementary Figure S7). We thus assigned a succinotype when the abundance of D was at least ten times higher than P, and *vice versa*, that is *r <* 0.1 or *r >* 0.9. Doing so, we were able to assign clear succinotypes to 7,653 (85.6%) of the retained amplicon samples and 11,442 (91.2%) of the retained shotgun samples.

**Figure 4:**
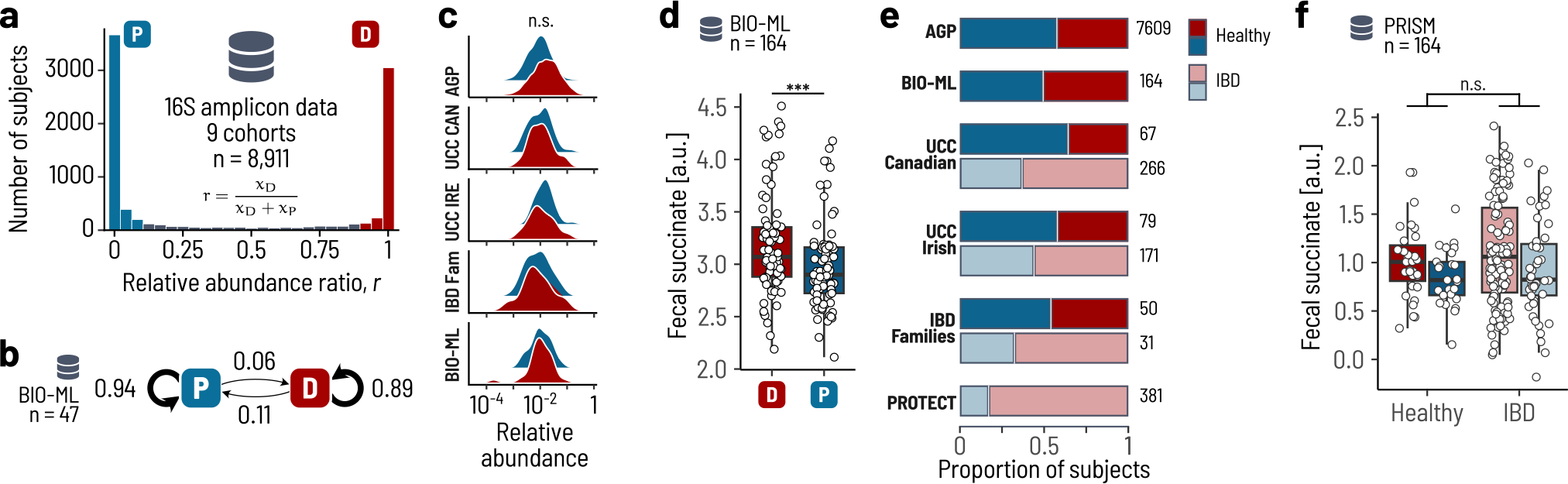
Clear succinotypes are found broadly and are associated with disease. **a**. The distribution of the relative abundance ratio of *Dialister* (D) compared to *Phascolarctobacterium*/*Phascolarctobacterium_A* (P) in fecal samples is strongly bimodal. The histogram encompasses 9 pooled cohorts of 16S amplicon data with a total of 8,911 individuals. **b**. Transition probabilities between succinotypes over time estimated individuals with repeat samples in the BIO-ML cohort. **c**. Distribution of relative abundances of D (red) and P (blue) in the fecal samples of healthy individuals. There is no significant difference between D and P across cohorts. **d**. Succinate concentrations in fecal samples from the BIO-ML cohort (Mann-Whitney U-test, *p* = 0.001619). **e**. Fraction of subjects assigned the D and P succinotype across cohorts and disease status. Dark colors are healthy subjects and light colors are IBD patients. **f**. Succinate concentrations in fecal samples from the PRISM cohort. There are no significant differences in fecal succinate between healthy subjects and IBD patients. AGP: American Gut Project; UCC CAN: UCC Canadian Cohort; UCC IRE: UCC Irish Cohort; IBD Fam: IBD Families Cohort; BIO-ML: Broad Institute-OpenBiome Microbiome Library.

We next asked whether the assignment of succinotypes to individuals remained stable over time using two of the cohorts. The Broad Institute-OpenBiome Microbiome Library (BIO-ML) and the Health Practitioners Follow-up Study (HPFS) comprise individuals that were sampled more than once. From these data, we computed the transition probabilities between succinotypes, that is, the fraction of times an individual changed its succino-types from D to P or *vice versa*. In BIO-ML, individuals retained the same succinotype in the subsequent fecal sample 88.9% and 94.4% of the time for D and P, respectively (Figure 4b), and the same was confirmed in the HPFS study (92% and 94%, Supplementary Figure S7). This demonstrates that individuals can be robustly classified into either a ‘D’ or ‘P’ succinotype based on their cross-sectional microbiome composition.

The relative abundance of the succinate utilizers did not differ between healthy individuals with different succinotypes. For each healthy subject, we computed the total relative abundance of succinate utilizers as the sum of relative abundances of D and P. The succinate utilizers had a mean relative abundance of 1.14%, with a 95% of the samples between 0.11% and 12.0% (Figure 4c). We tested for a difference in relative abundance of the succinate utilizers between individuals that were assigned to either succinotype. To accounts for potential variability between datasets, we used a mixed-effects model with random slopes and intercepts for the succinotype across cohorts. There was no consistent significant effect of succinotype on the log relative abundance (*p* = 0.503). This implies that the size of the ‘succinate utilization niche’ is conserved and hence does not depend on the identity of the taxon that occupies the niche.

We hypothesized that the lower rate of succinate consumption by *Dialister* as compared to *Phascolarctobacterium* would imply that the net concentration of succinate is higher in D-type as compared to P-type individuals. To test this, we used the metabolomic data from the BIO-ML cohort. Fecal succinate concentrations were significantly higher in D-types compared to P-types (Mann-Whitney U-test, *p* = 0.001619; Figure 4d), and there was no significant effect of the relative abundance of succinate utilizers in a linear model including both succinotype and relative abundance (*p* = 0.559, Supplementary Figure S8). These results support the notion that the niche size of succinate consumption does not differ between succinotypes, but that the slower removal of succinate by *Dialister* manifests as a higher net succinate concentration.

Given these increased levels of intestinal succinate and the reported role of succinate in inflammation^3^, we hypothesized that a *Dialister* succinotype would be more prone to intestinal inflammation—and possibly at higher risk of inflammatory bowel disease (IBD). We tested for an effect of succinotype on IBD status using all cohorts in a logistic regression on succinotype with cohort as a random effect.

IBD patients were consistently more likely to be D-types than P-types (Figure 4e). The odds ratio of IBD *versus* healthy was 2.2 times lower for P-types compared to D-types (logit regression, *β* = *−*0.805, CI = [*−*1.16, *−*0.449], *p* = 9.27 *·* 10^*−*6^). We thus asked whether succinate concentrations are disproportionately higher in D-type IBD patients than P-types using the metabolomic data from the PRISM cohort^30^, for which taxonomic composition based on shotgun metagenomic data paired with metabolomic data were available^31^. Consistent with the data from BIO-ML, fecal succinate concentrations were higher in D-types than in P-types, but the difference was not significantly different from that in healthy individuals (Figure 4f; two-way ANOVA, *F*_IBD_(1, 217) = 2.32, *p*_IBD_ = 0.12, *F*_Stype_(1, 217) = 6.29, *p*_Stype_ = 0.013). Overall, this suggests that the *Dialister* succinotype contributes in some manner to IBD pathogenesis, likely as the result of its slower consumption of succinate.

Finally, to check to which extent the association of succintypes with disease was specific to IBD, we looked at the succinotype distributions in a set of other diseases that were part of the curatedMetagenomicDatasets package. Two diseases, colorectal cancer (CRC) and atherosclerotic cardiovascular disease (ACVD), were associated with the P succinotype (Supplementary Figure S9). However, none of the other tested diseases were significantly associated with the D succinotype, indeed suggesting a specific role of high *Dialister* or low *Phascolarctobacterium* in IBD.

## Discussion

The goal of this study was to map out which human intestinal bacteria are the main consumers of succinate—a metabolite that is at a key crossroad of human and microbiome metabolism. Our multi-faceted analysis of fecal microbiota and bacterial isolates identified two bacterial taxonomic groups as key succinate consumers in human intestinal microbiomes, *Phascolarctobacterium* and *Dialister*. These two taxa are highly mutually exclusive in fecal microbiomes of western populations, allowing for the clear classification of individuals into P and D ‘succinotypes’. These succinotypes differ with respect to their succinate consumption rate, with the D types consuming succinate more slowly than the P-succinotypes, and this translates to higher fecal concentrations of succinate. Finally, while the prevalence of succinotypes in healthy populations is rather balanced, IBD patients are significantly more likely to have a D-succinotype than a P-succinotype. This provides evidence for an imbalance between host and microbiota succinate metabolism in intestinal inflammatory diseases like IBD.

The discovery that only two taxonomic groups occupy the succinate niche in human intestinal microbiota is strikingly simple, and somewhat unexpected given the wider diversity of intestinal bacteria that have been reported to consume succinate. These include, for example, *Clostridioides difficile* and *Clostridium kluyveri* that convert succinate to butyrate^32,33^, or *Veillonella parvula* that decarboxylates succinate during lactate consumption^17^. However, these bacteria do not *per se* use succinate as a growth substrate, with the former two regenerating NAD+ and the latter increasing growth yield from lactate but not able to grow on succinate alone. This might be similar to how *Flavonifractor plautii* uses succinate in our data, with a comparatively low *per capita* consumption rate. In contrast, *Phascolarctobacterium* and *Dialister* generate energy for growth from the decarboxylation of succinate. Yet, because the energetic yield of converting succinate to propionate is low^34^, there is limited selection pressure for bacteria to specialize on succinate consumption. This is further illustrated by those bacteria that encode the full succinate pathway, like *Bacteroides* sp., but mostly favor cutting short and excreting succinate instead of fully running the pathway to produce propionate. Using succinate as a growth substrate has been observed for other taxa, including *Propionigenium modestum*^35^ and *Peptostreptococcus* sp.^34^, but given that (a) *Phascolarctobacterium* and *Dialister* are *de facto* fully mutually exclusive and (b) the prevalence of either is close to 100% in humans, we posit that these two taxonomic groups have specialized specifically on succinate consumption in the human intestine. What the evolutionary process was that resulted in this arrangement remains to be investigated.

To what degree the observed consumption rate differences translate to what happens *in situ* needs to be carefully evaluated. Our fecal enrichment approach was designed to probe bacterial competitive ability in the relevant background of the full complex microbiota, but by design makes specific choices of the growth environment and also neglects the host. Indeed, the consumption of succinate is dependent on a number of cofactors, in particular vitamin B_12_^36^ and sodium^34,35^, and the specific local concentrations of these and other cofactors need not be the same in our lab setup and in humans. We do observe a decrease in succinate in all tested fecal samples, suggesting that our lab setup does allow for succinate consumption to occur. A more conservative interpretation of the difference in consumption rates is that *Phascolarctobacterium* has a more robust succinate consumption system than *Dialister* and is thus less dependent on externally supplied cofactors and conditions. However, the observed higher fecal succinate concentrations in D-succinotype individuals compared to P-succinotypes despite equal population sizes does provide independent evidence for an *in vivo* difference in consumption rate.

Our results provide a rare mechanistic link between microbiome composition and function, and are a basis on which to functionally interpret compositional dysbiosis in disease. Differential abundances of both *Phascolarctobacterium* and *Dialister* have been independently reported as associated with IBD, but have mostly not been interpreted together. The first analyses of the OSCCAR and PRISM cohorts already reported *Phascolarctobacterium* as one of the two only genera that were significantly reduced in UC and CD patients^37^ with no specific mention of *Dialister*, an observation independently confirmed in other cohorts^38,39^. Subsequent studies also reported concomitant decreased abundances of *Phascolarctobacterium* and increased *Dialister* between IBD patients and healthy controls^39,40^. While these studies did point into the direction of a role of intestinal succinate in IBD, our classification into functionally different succinotypes now allows for a mechanistic interpretation of these consistent signals.

Attempting to classify human microbiota into different types is not new, the most prominent example being the ‘enterotypes’^41,42^. Our succinotype classification differs from the enterotype classification in an important way: succinotypes are primarily informed by a functional phenotype, while enterotypes are informed by statistical differentiation of microbiome composition. While such statistical classification can be useful to reduce complex compositional differences into a simple grouping, there is *per se* no direct interpretation of what this grouping means. In constrast, the two succinotypes we describe here are based on succinate consumption rate and thus can be directly interpreted.

Finally, the succinotype classification allows for a robust stratification of patients based on a clinically relevant feature. Clinical parameters can then be tested for association with disease. In our analysis, for example, fecal succinate concentrations are higher in IBD patients compared to healthy controls, but this difference is better explained by succinotype rather than disease status. Such biomarker-based patient stratification is a powerful tool to inform treatment decisions and ultimately improve treatment outcomes.

## Acknowledgements

We thank Leandra Knecht for support with experimental work, Alice de Wouters and Shaul Pollak for feedback and discussions, and Jill Marie O’Sullivan for uncovering additional studies reporting differential abundances of P and D. We thank the AVINA Foundation for financial support.

## Author Contributions

L.A., C.M., and J.Z. performed experiments. L.A., T.d.W., and G.E.L. planned the execution of the project. L.A., P.R.v.B., and G.E.L. analyzed the data. L.A., T.d.W., and G.E.L. conceived the project. L.A., C.L., T.d.W., and G.E.L. interpreted the results. G.E.L. and L.A. wrote the first draft of the manuscript. All authors reviewed the final manuscript and provided feedback.

## Data availability

The raw data is available at Figshare during peer review and will be made public upon publication.

## Methods

### Stool collection

The research project was approved by the Ethic Committee of the Canton of Zurich (2017-01290). Fresh fecal samples were donated from 13 healthy individuals with no history of antibiotic use, intestinal infections, or severe diarrhea during the three months prior to making the donation. The donors did not take immunosuppressive drugs, blood thinners, or medication affecting the bowel passage or digestion. Fecal samples were anaerobically transported in an airtight container together with an Oxoid™ AnaeroGen™ 2.5 L sachet (Thermo Fisher Diagnostics AG, Pratteln, Switzerland) and processed within three hours after defecation. Stool consistency was evaluated optically according to the Bristol Stool Scale^43^ and samples within the defined range of a healthy stool, notably with a score between 3-5, were accepted.

### Preparation of anaerobic culture media

The growth medium was adapted from the M2GSC medium^44^ for the *in vitro* enrichments of complex cultures as in Anthamatten *et al*.^22^, and additionally also from YCFA^45^ for single cultures. Briefly, in contrast to the common M2GSC and YCFA media, our basal medium did not contain any specific carbon sources, i.e. no glucose, starch, or cellobiose and had reduced concentrations of amicase (1 g*/*L), yeast extract (1.25 g*/*L), and meat extract (0.5 g*/*L). All medium ingredients except sodium bicarbonate and L-cysteine HCl were dissolved in an Erlenmeyer flask and the pH was adjusted to pH 7 using 5 mM sodium hydroxide. The media were boiled for 15 min under constant moderate stirring to removal oxygen and using a Liebig condenser to prevent vaporization of ingredients. After boiling, the media were constantly flushed with CO_2_. Sodium bicarbonate and L-cysteine hydrochloride monohydrate were added when the media had cooled to 55 ^*◦*^C for further reduction of residual oxygen for 10 min. Aliquots of 8 mL of medium were filled into Hungate tubes under constant flushing with CO_2_, and Hungate tubes were sealed with butyl rubber stoppers and screw caps (Millan SA, Geneva, Switzerland). The media were subsequently sterilized by autoclaving and stored at room temperature.

### Stool processing

The fecal samples were transferred into an anaerobic chamber (10% CO_2_, 5% H_2_, and 85% N_2_) (Coy Laboratories, Ann Arbor, MI, USA). One gram of fecal sample measured with a sterile plastic spoon (VWR International, Dietikon, Switzerland) was suspended in 9 mL of anaerobic dilution solution (ADS). The dilution step was repeated and 1 mL of the resulting 100-fold dilution was transferred into 9 mL of ADS in a sterile Hungate tube. Serial dilutions down to 10^*−*11^ were continued outside of the anaerobic chamber under sterile anaerobic conditions using the Hungate technique to determine the total viable cells after transport using the most probably number method.

### Batch enrichments in succinate-rich conditions

Anaerobic *in vitro* enrichments were performed in Hungate tubes sealed with butyl rubber stoppers and screw caps (Millan SA, Geneva, Switzerland). For each enrichment, 0.3 mL of the 10^*−*8^ fecal sample dilution was inoculated into 8 mL of cultivation medium in Hungate tubes. Enrichments were performed for each of the 13 studied microbiota in three replicate cultures in a basal medium supplemented with 30 mM of disodium succinate and in an non-supplemented control condition. The media were buffered at an initial pH of 6.5. All cultures were incubated at 37 ^*◦*^C for up to seven days. The optical densities were measured at 600 nm directly in the Hungate tubes with a WPA CO 8000 Cell Density Meter (Biochrom Ltd, Cambridge, England).

### Microbial metabolite analysis

Succinate concentrations were measured by HPLC analysis. Samples were prepared from 1 mL of bacterial culture centrifuged at 14 000 g for 10 min at 4 ^*◦*^C. The supernatant was filtered into 2 mL short thread vials with crimp caps (VWR International GmbH, Schlieren, Switzerland) using non-sterile 0.2 µm regenerated cellulose membrane filters (Phenomenex Inc., Aschaffenburg, Germany). A volume of 40 µL of sample was injected into the HPLC with a flow rate of 0.6 mL*/*min at a constant column temperature of 80 ^*◦*^C and using a mixture of H_2_SO_4_ (10 mM) and Na-azide (0.05 g*/*L) as eluent. Analyses were performed with a Hitachi Chromaster 5450 RI-Detector (VWR International GmbH, Schlieren, Switzerland) using a Rezex ROA-Organic Acid (4 %) precolumn connected to a Rezex ROA-Organic Acid (8 %) column, equipped with a Security Guard Carbo-H cartridge (4 *×* 3 mm). Metabolite concentrations were determined using external standards (all purchased from Sigma-Aldrich, Buchs, Switzerland) via comparison of the retention times. Peaks were integrated using the EZChromElite software (Version V3.3.2.SP2, Hitachi High Tech Science Corporation).

### DNA extractions and compositional profiling by 16S metagenomic sequencing

The total genomic DNA was extracted from 200 mg of stool or from pellets of 1 mL culture (centrifuged at 14 000 g for 10 min at 4 ^*◦*^C), using the Maxwell®RSC PureFood GMO and Authentication Kit. The quality of all DNA extracts was confirmed on a Tris-Acetate-EDTA (TAE)-1.5 % agarose gel. The total DNA concentration after extraction was quantified using the Qubit®dsDNA HS Assay kit. We performed amplicon sequencing of the 16S rRNA V3-V4 region on Illumina MiSeq using the primer combination 341F (5’-CCTACGGGNBGCASCAG-3’) and 806bR (5’-GGACTACNVGGGTWTCTAAT-3’). Library preparation and sequencing was performed by StarSEQ GmbH (Mainz, Germany) with 25 % PhiX. Amplicon Sequence Variants (ASVs) were inferred using Dada2 v1.18.0. Forward and reverse reads shorter than 250 and 210 were filtered. The maximum number of expected errors was set to 4 and 5 for forward and reverse reads, respectively. Inference was performed in ‘pseudo pool’ mode. Read pairs were merged with a minimum overlap of 20, and chimeras were removed using the ‘consensus’ method. Taxonomic annotation was done with the RDP classifier of Dada2 using the GTDB r95 database^46^.

### Succinate-specific enrichment

We first determined the genus-level composition of each of the succinate supplemented (SU) and non-supplemented (SU–) enrichment cultures based on 16S amplicon sequencing. To this end, we computed the regularized genus-level relative abundance by grouping all ASVs that were taxonomically classified as the same genus, and normalizing the genus-specific read counts by the total read count in the sample after adding a pseudo-count of 1 if a specific genus had at least one count in the fecal sample or any derived enrichment culture. Note, that in a Bayesian interpretation the pseudo-count of 1 corresponds to a flat Dirichlet prior in a Dirichlet-Multinomial model for read counts. To account for unspecific growth, we determined maximum regularized relative abundance of a genus in the SU– enrichments. We defined the succinate-specific enrichment as the difference in regularized relative abundance of the genus in the SU cultures minus the unspecific growth.

We tested for associations of genera with succinate consumption categories of the fecal microbiota using a statistical approach. We first filtered for those genera for which the maximum succinate enrichment across all enrichments was at least 0.01, yielding 35 out of a total of 292 genera. With these 35 genera, we then performed a joint linear regression of the enrichment score on the interaction of consumption category, day, and genus with no intercept. This tests, for each combination genus, category, and day, whether the succinate enrichment is statistically different from zero. We then selected those genera with *p <* 0.05 and an estimated mean larger than zero. Using this approach we identified the four genera *Phascolarctobacterium, Phascolarctobacterium_A, Dialister*, and *Flavonifractor* (Supplementary Figure S1).

### Identification of the mmdA gene and succinate pathway

The goal was to broadly identify putative succinate consumers based on the similarity to the mmdA gene of the *Veillonella parvula* succinate-to-propionate cluster (UniProt Q57079). We first looked for putative hits in all translated genomes from GTDB release 214 with phmmer (HMMER 3.3) and filtered for those with an *e*-value below 10^*−*150^, resulting in 1,643 distinct genomes. For each genus, we kept only the hit with the lowest *e*-value, leaving 281 distinct genera. We then performed multiple sequence alignment with MAFFT^47^ 7.453 including the human PCCB gene to use as a phylogenetic outgroup. Finally, we reconstructed the phylogeny of mmdA genes using RAxML-NG^48^ 1.2.0 with a JTT+G model, 10 maximum parsimony starting trees, and 200 bootstraps. For representative genomes of the four succinate utilizers (GCA_000160055.1, GCA_003945365.1, GCA_010508875.1, GCA_023497905.1), we identified the succinate-to-propionate gene cluster using gutSMASH^49^.

### Testing of isolates for succinate consumption

Strains were pre-cultured in M2GSC medium, with the exception of *Akkermansia muciniphila* DSMZ 22959 that was pre-cultured in M2-based medium supplemented with 3 g*/*L of type II mucin (Sigma-Aldrich Chemie GmbH, Buchs, Switzerland), and *Veillonella parvula* DSMZ 2008 that was pre-cultured in M2-based medium supplemented with 60 mM of DL-lactic acid 90% (Sigma-Aldrich Chemie GmbH, Buchs, Switzerland). To test for succinate consumption, we inoculated 0.1 mL of 48 h pre-cultures into 8 mL of M2-based medium supplemented with 30 mM or 80 mM of succinate, and quantified the succinate concentrations after two and seven days.

### Cultures for succinate consumption kinetics

The strains to be tested were pre-cultured in YCFA medium supplemented with 80 mM of succinate. To standardize the starting cell densities, we quantified the cell concentrations of the pre-cultures by flow cytometry using live/dead staining. A double staining assay with the two nucleic acid dyes SYBR Green (SG) and propidium iodide (PI) was used to differentiate between cells with intact (viable) and damaged (dead) cytoplasmic membranes^50^. We aimed for a starting cell density of 1 *×* 10^7^ cells*/*mL, if necessary, pre-cultures were diluted accordingly in anaerobic dilution solution.

Cultures were performed in 8 mL of YCFA medium supplemented with 80 mM of succinate. We selected sampling times for the different strains based on preliminary growth test in order to target the time window where most of the succinate consumption occurs: 1 h intervals during 25 h for *Phascolarctobacterium*; 2 h intervals during 52 h for *Flavonifractor*; and 4 h intervals during 100 h for *Dialister*. Separate cultures were inoculated for each time point to avoid effects from repeated sampling, and full duplicates using separate pre-cultures.

### Estimation of the *per capita* succinate consumption rate

We modelled succinate uptake following Monod,

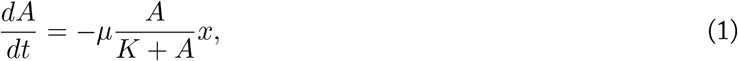

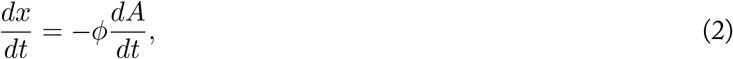

where *A*(*t*) is the concentration of substrate and *x*(*t*) is the microbial density at time *t. µ* is the maximum *per capita* succinate uptake rate, and *ϕ* is the growth yield per unit succinate taken up.

We then estimate the parameters *µ, ϕ*, and *K* using Bayesian Markov Chain Monte Carlo (MCMC) using Stan^51,52^. For each sample of the parameters, we first numerically solve the system of ordinary differential equations (1) and (2) with *A*(0) = *A*_0_ and *x*(0) = *x*_0_. We then compute the log-likelihood of the observed data using the following hierarchical model,

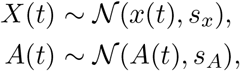

where *N* signifies the normal distribution, with the parameters sampled from the following prior distributions unless otherwise specified,

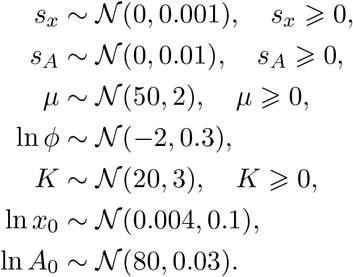

We use the Runge-Kutta (4,5) method as implemented in CmdStan to solve the system of ODEs. Summary statistics for the sampled parameters and convergence diagnostics are listed in Supplementary Text S1.

To account for bacterial growth on a first (unobserved) preferential substrate, we modified equations (1) and (2) to include a relative allocation into the uptake of a preferential substrate, *B*, and a secondary substrate, *A*. For simplicity, we use the affinity fraction, *B/*(*K*_*B*_ + *B*), from the Monod equations as the switch between substrates,

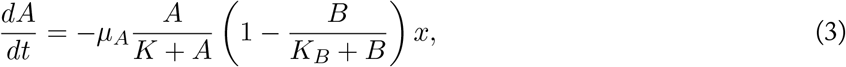

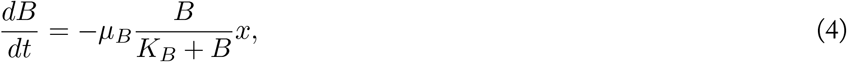

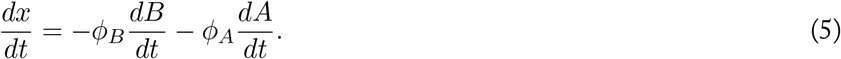

We then estimated the parameters in an analogous MCMC approach as the simpler model, with the additional or modified priors,

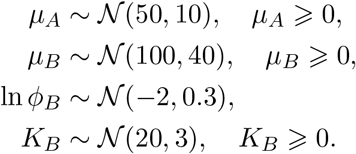

Because the concentration of the substrate *B* is not observed, we cannot estimate the values of *B*_0_, which we set to *B*_0_ = 100 without loss of generality. This implies that the value of *B* and thus also *µ*_*B*_, *ϕ*_*B*_, and *K*_*B*_ are in arbitrary units.

### Other amplicon and shotgun metagenomic data

For the American Gut Project^24^, we downloaded the sOTU tables and corresponding DNA sequences from figshare and performed a new taxonomic assignment with the GTDB r95 database and the RDP classifier from Dada2. For the PROTECT^27^ and UCC^25^ data, we downloaded the raw sequencing reads from NCBI PRJNA436359 and PRJNA414072, and inferred ASVs and performed taxonomic assignment using the same pipeline as for the enrichment data. Additionally, we fetched those datasets from the ‘gut microbiome-metabolome dataset colletion’^31^ that were based on 16S amplicon data^23,26,53–55^, and directly used the ASV counts and GTDB taxonomic assignments.

For the shotgun metagenomic data, we fetched all datasets from the curatedMetagenomicData^28^ for which relative abundance were computed and used the provided taxonomic assignments. Additionally, we also fetched the compositional and metabolomic data for PRISM^30^ cohort from the ‘gut microbiome-metabolome dataset colletion’.

---

**Figure S1:**
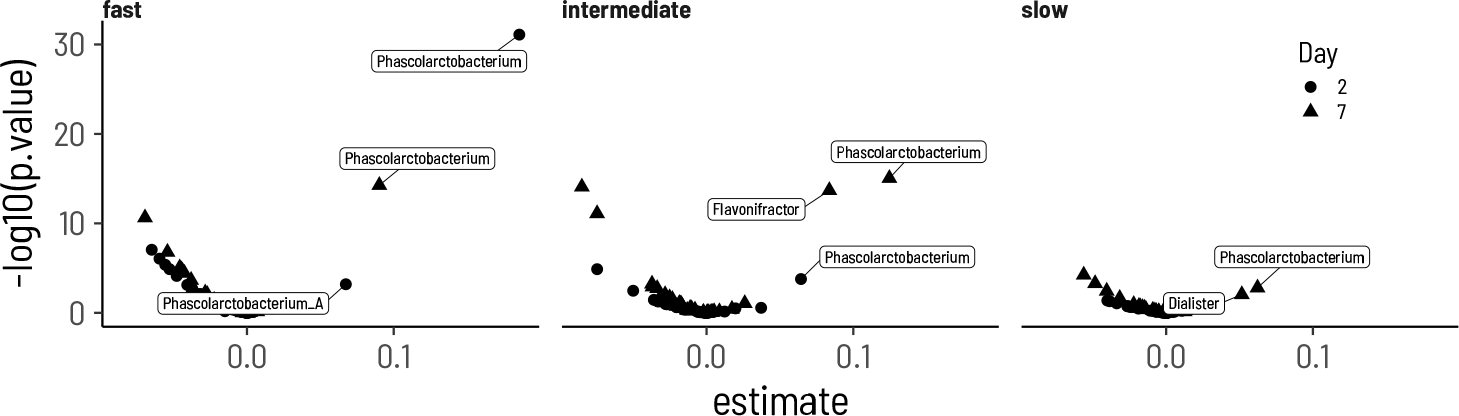
Regression coefficients of the genus enrichment and the consumption category. The estimates are obtained from a linear regression of the difference in relative abundance between the SU+ and SU-condition and the rate category split by day. Estimates with a value greater than zero and p < 0.05 are labelled.

**Figure S2:**
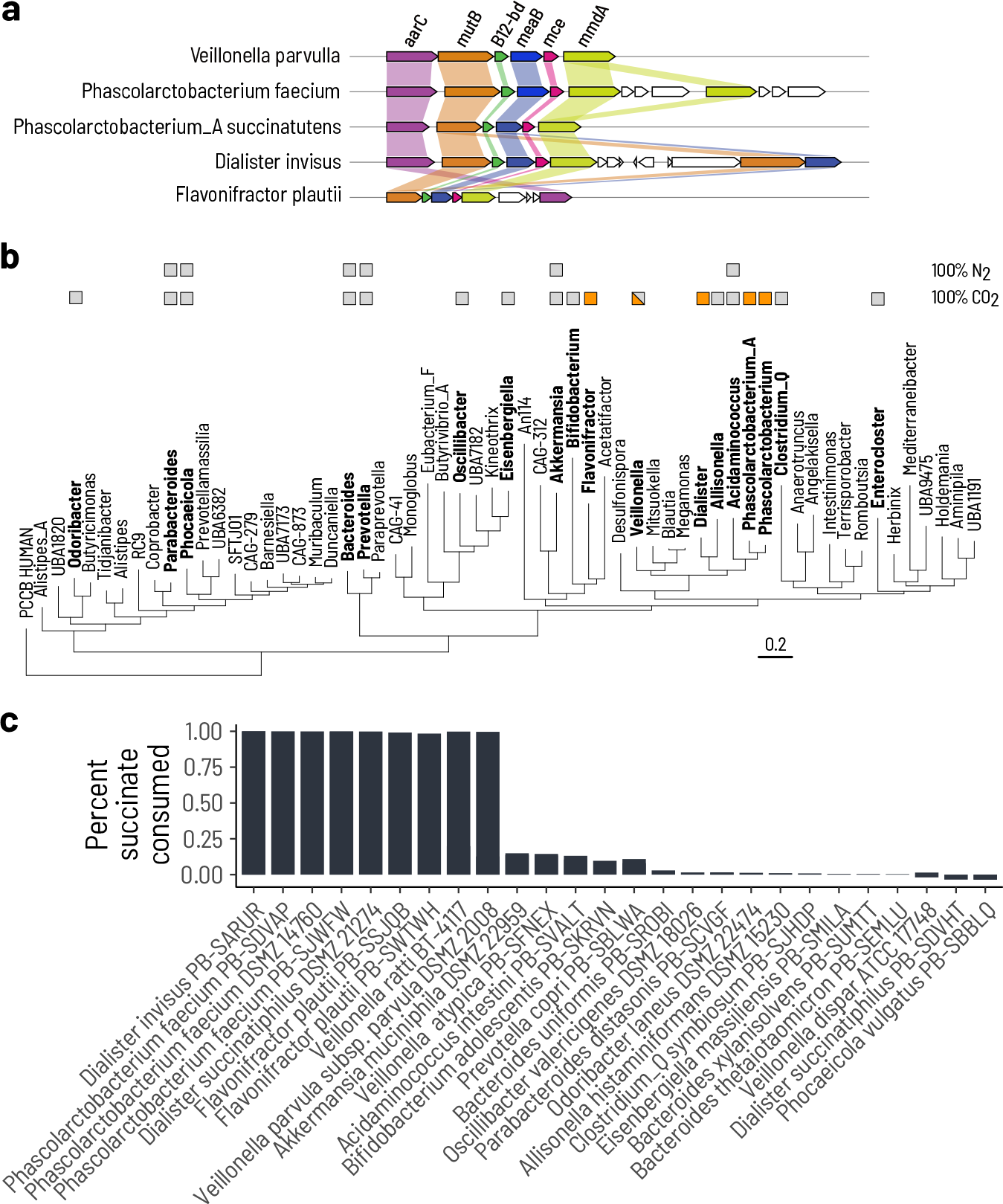
Extracellular succinate consumption is restricted to few bacterial taxa. **a**. The gene cluster associated with succinate to propionate conversion occurs in all succinate consumers. **b**. Phylogenetic tree based on the mmdA gene. Isolates that were screened for succinate consumption are indicated in bold. Succinate consumption is indicated with an orange box, and non-consumption with a grey box. Partial consumption is indicated by both colors. Consumption was also tested under a 100% N_2_ atmosphere for a subset of isolates. **c**. Maximum percentage of supplemented succinate that was consumed by each tested isolate.

**Figure S3:**
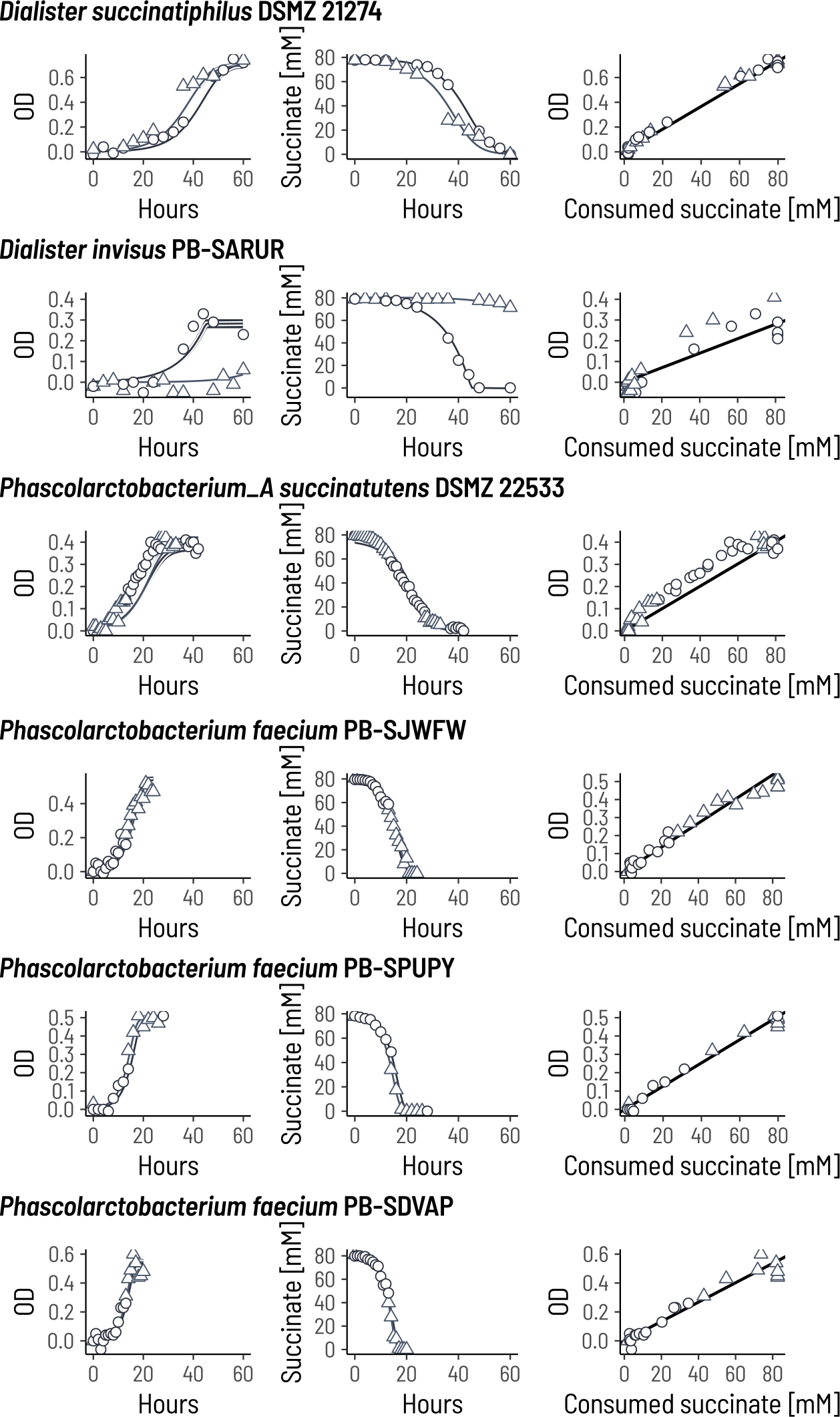
A mathematical model of substrate uptake fits well to the genera *Phascolarctobacterium* and *Dialister*. Each point is an independent culture destructively sampled at the specified time to measure optical density (OD) and succinate concentration. Two independent inoculum cultures were prepared for each strain (circles and triangles). The lines show the posterior median curve across all samples from the posterior distribution. The shaded area (where visible) show the 95% HPD interval across the posterior samples. We population size of the inoculum cultures were estimated independently, resulting in two lines. The slope of the line in the rightmost panels is the posterior mean of the OD yield per unit succinate parameter, *ϕ*.

**Figure S4:**
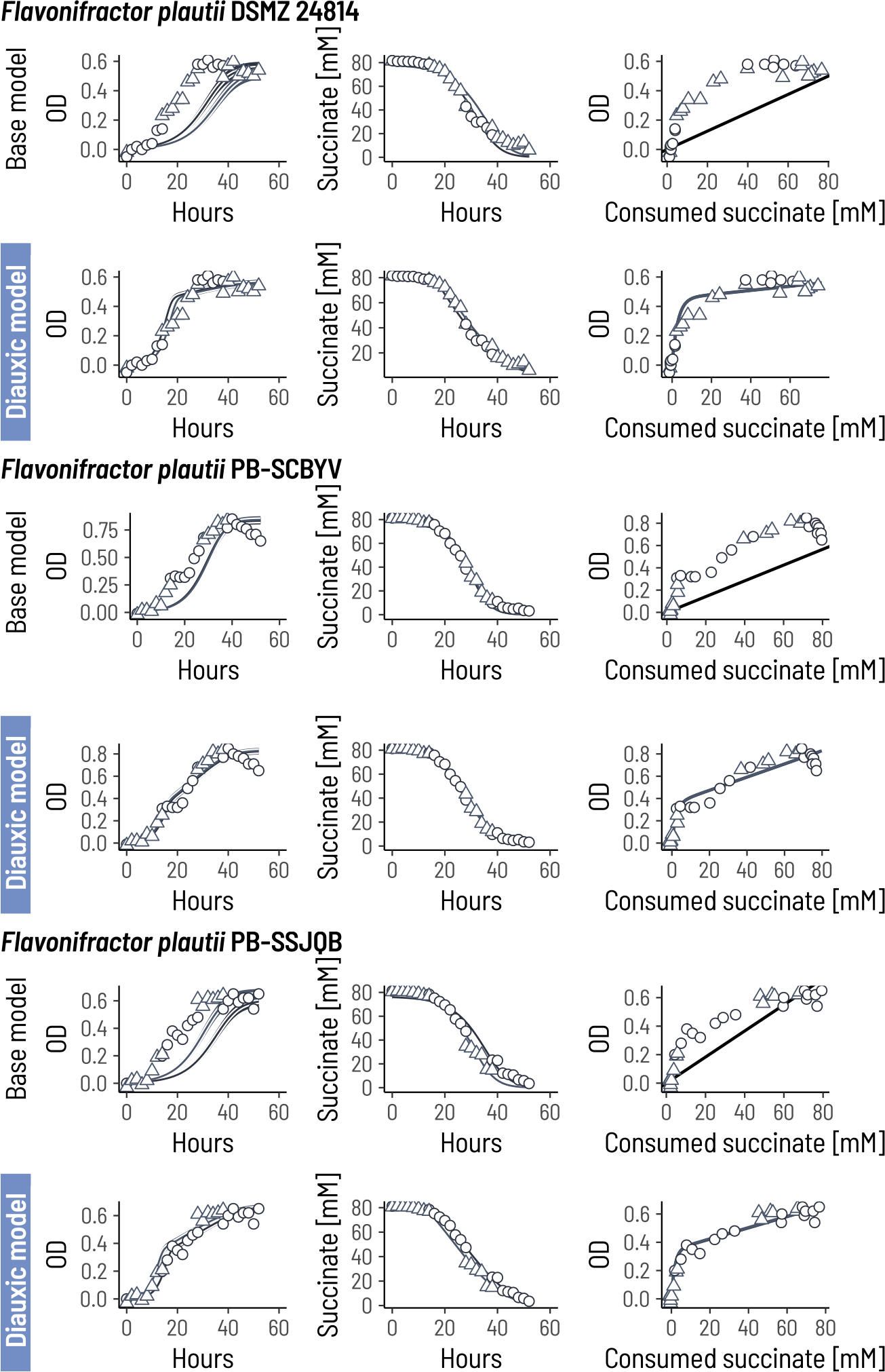
A mathematical model with diauxie is a better fit for the genus *Flavonifractor*. Each point is an independent culture destructively sampled at the specified time to measure optical density (OD) and succinate concentration. Two independent inoculum cultures were prepared for each strain (circles and triangles). The lines show the posterior median curve across all samples from the posterior distribution. The shaded area (where visible) show the 95% HPD interval across the posterior samples. We population size of the inoculum cultures were estimated independently, resulting in two lines. For the base model, the slope of the line in the rightmost panels is the posterior mean of the OD yield per unit succinate parameter, *ϕ*. For the diauxic model, the lines show the median across the samples from the posterior.

**Figure S5:**
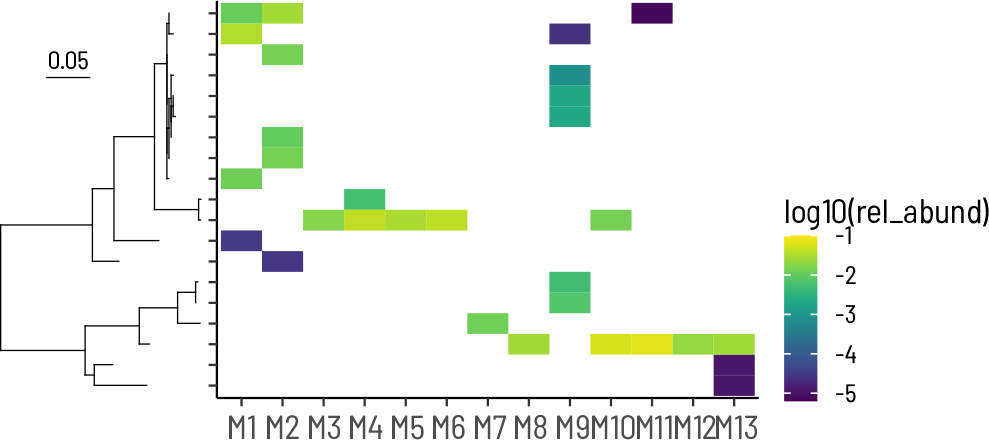
Relative abundances of all ASVs classified as *Phascolarctobacterium, Phascolarctobacterium_A*, or *Dialister*.

**Figure S6:**
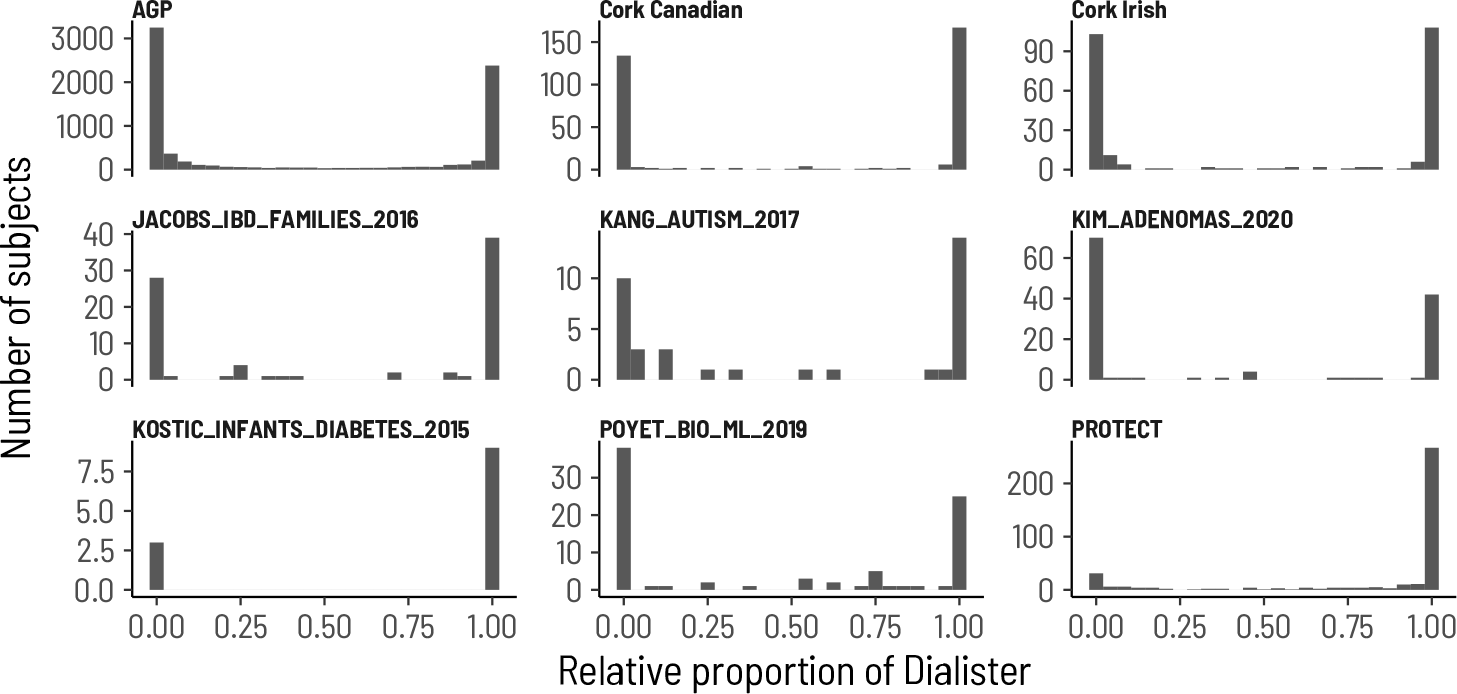
Distribution of the succinotype ratio across cohorts. The ratio of abundances *r* = *xD/*(*xD* + *xP*) is strongly bimodal across all cohorts.

**Figure S7:**
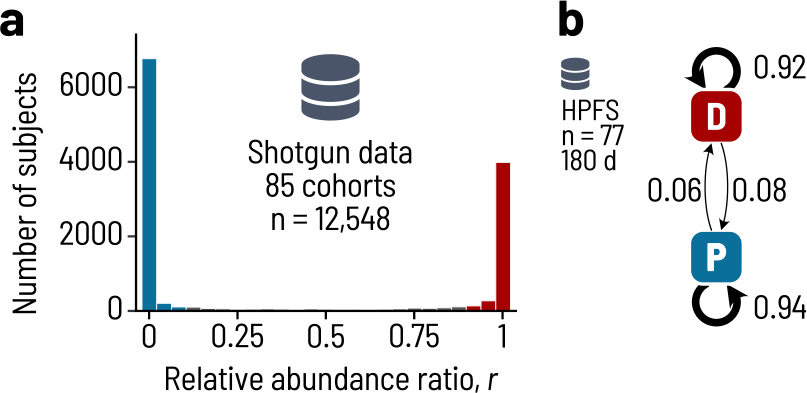
Succintypes in cohorts with shotgun metagenomic data. **a**. The ratio of abundances *r* = *xD/*(*xD* + *xP*) is strongly bimodal across all cohorts. **b**. Probability of transitioning from one succinotype to another in the Health Professionals Follow-up Study (HPFS).

**Figure S8:**
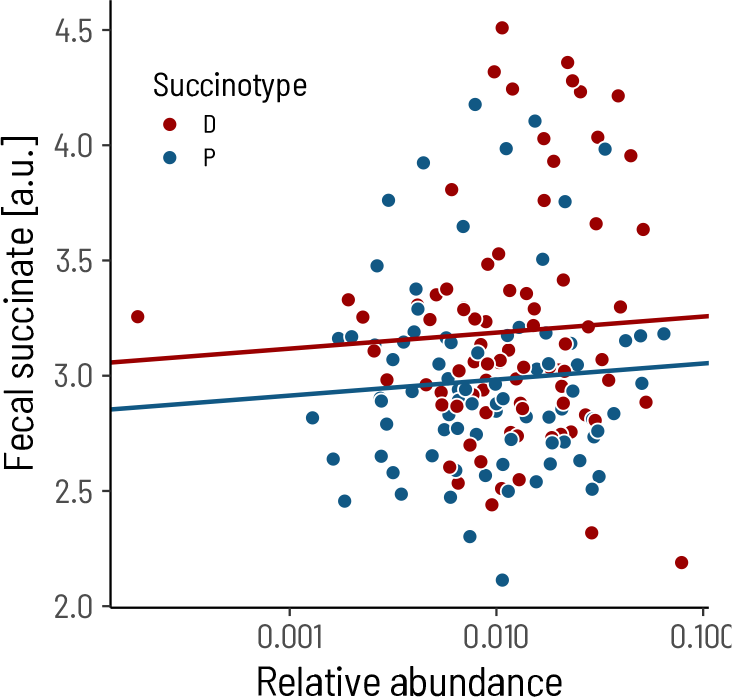
Fecal succinate concentrations do not depend on the relative abundance of the succinate consumers. BIO-ML cohort, n = 146. The slopes are estiamted from a linear model with succinotype and log10 relative abundance as fixed effects.

**Figure S9:**
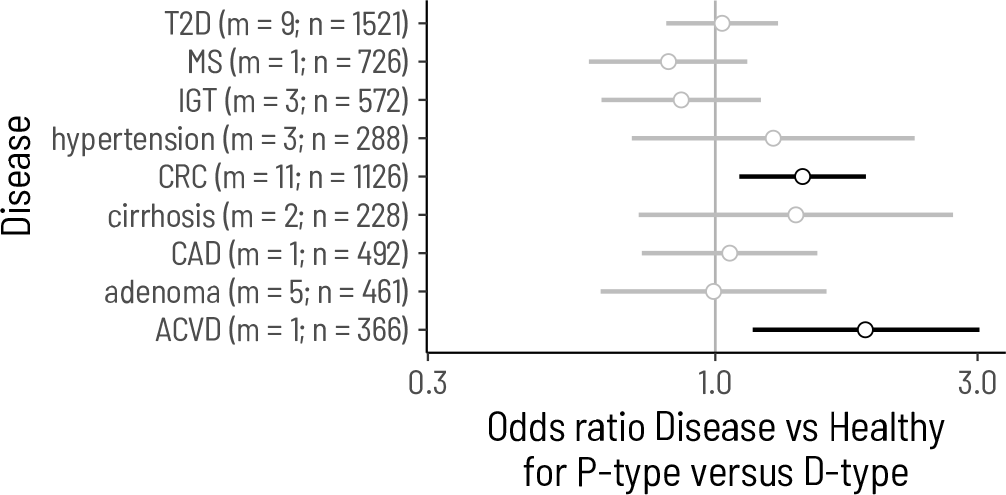
Association of succinotypes with diseases other than IBD. Estimates are from logistic regressions of disease *versus* healthy on succinotype, including the study as a random effect if there were more than 1 study. m, number of studies; n, number of subjects. T2D, type 2 diabetes; MS, multiple sclerosis; IGT, impaired glucose tolerance; CRC, colorectal cancer; CAD, cardiovascular disease; ACVD, atherosclerotic cardiovascular disease.

## Supplementary Text S1

The following pages show the summary statistics and convergence diagnostics for the MCMC sampling.

## MCMC diagnostics

### Model 1

#### DSMZ-109768

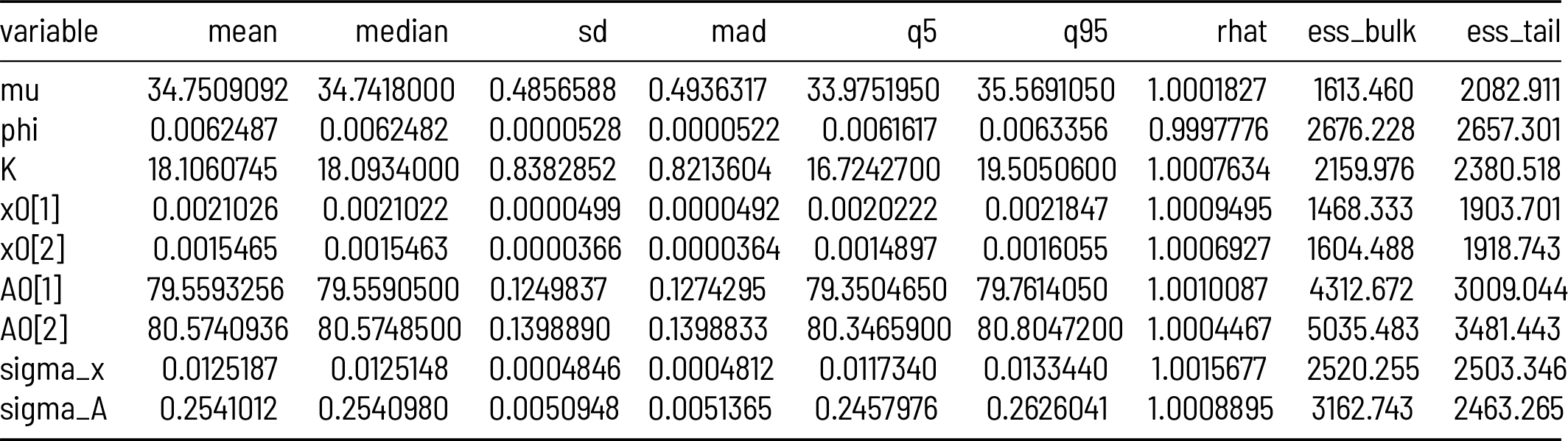

| variable | mean | median | sd | mad | q5 | q95 | rhat | ess_bulk | ess_tail |
| --- | --- | --- | --- | --- | --- | --- | --- | --- | --- |
| mu | 34.7509092 | 34.7418000 | 0.4856588 | 0.4936317 | 33.9751950 | 35.5691050 | 1.0001827 | 1613.460 | 2082.911 |
| phi | 0.0062487 | 0.0062482 | 0.0000528 | 0.0000522 | 0.0061617 | 0.0063356 | 0.9997776 | 2676.228 | 2657.301 |
| K | 18.1060745 | 18.0934000 | 0.8382852 | 0.8213604 | 16.7242700 | 19.5050600 | 1.0007634 | 2159.976 | 2380.518 |
| x0[1] | 0.0021026 | 0.0021022 | 0.0000499 | 0.0000492 | 0.0020222 | 0.0021847 | 1.0009495 | 1468.333 | 1903.701 |
| x0[2] | 0.0015465 | 0.0015463 | 0.0000366 | 0.0000364 | 0.0014897 | 0.0016055 | 1.0006927 | 1604.488 | 1918.743 |
| A0[1] | 79.5593256 | 79.5590500 | 0.1249837 | 0.1274295 | 79.3504650 | 79.7614050 | 1.0010087 | 4312.672 | 3009.044 |
| A0[2] | 80.5740936 | 80.5748500 | 0.1398890 | 0.1398833 | 80.3465900 | 80.8047200 | 1.0004467 | 5035.483 | 3481.443 |
| sigma_x | 0.0125187 | 0.0125148 | 0.0004846 | 0.0004812 | 0.0117340 | 0.0133440 | 1.0015677 | 2520.255 | 2503.346 |
| sigma_A | 0.2541012 | 0.2540980 | 0.0050948 | 0.0051365 | 0.2457976 | 0.2626041 | 1.0008895 | 3162.743 | 2463.265 |

**DSMZ-14760**

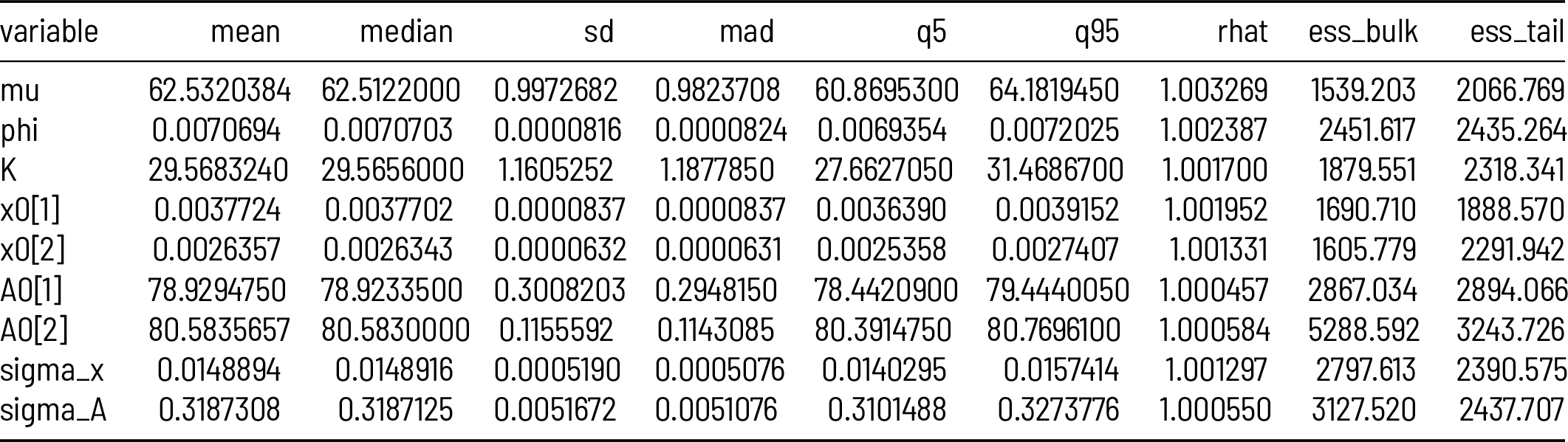

| variable | mean | median | sd | mad | q5 | q95 | rhat | ess_bulk | ess_tail |
| --- | --- | --- | --- | --- | --- | --- | --- | --- | --- |
| mu | 62.5320384 | 62.5122000 | 0.9972682 | 0.9823708 | 60.8695300 | 64.1819450 | 1.003269 | 1539.203 | 2066.769 |
| phi | 0.0070694 | 0.0070703 | 0.0000816 | 0.0000824 | 0.0069354 | 0.0072025 | 1.002387 | 2451.617 | 2435.264 |
| K | 29.5683240 | 29.5656000 | 1.1605252 | 1.1877850 | 27.6627050 | 31.4686700 | 1.001700 | 1879.551 | 2318.341 |
| x0[1] | 0.0037724 | 0.0037702 | 0.0000837 | 0.0000837 | 0.0036390 | 0.0039152 | 1.001952 | 1690.710 | 1888.570 |
| x0[2] | 0.0026357 | 0.0026343 | 0.0000632 | 0.0000631 | 0.0025358 | 0.0027407 | 1.001331 | 1605.779 | 2291.942 |
| A0[1] | 78.9294750 | 78.9233500 | 0.3008203 | 0.2948150 | 78.4420900 | 79.4440050 | 1.000457 | 2867.034 | 2894.066 |
| A0[2] | 80.5835657 | 80.5830000 | 0.1155592 | 0.1143085 | 80.3914750 | 80.7696100 | 1.000584 | 5288.592 | 3243.726 |
| sigma_x | 0.0148894 | 0.0148916 | 0.0005190 | 0.0005076 | 0.0140295 | 0.0157414 | 1.001297 | 2797.613 | 2390.575 |
| sigma_A | 0.3187308 | 0.3187125 | 0.0051672 | 0.0051076 | 0.3101488 | 0.3273776 | 1.000550 | 3127.520 | 2437.707 |

**DSMZ-21274**

| variable | mean | median | sd | mad | q5 | q95 | rhat | ess_bulk | ess_tail |
| --- | --- | --- | --- | --- | --- | --- | --- | --- | --- |
| mu | 29.3693966 | 29.3803500 | 0.5140788 | 0.5223200 | 28.5100800 | 30.2277250 | 1.001576 | 1871.567 | 2269.861 |
| phi | 0.0091525 | 0.0091537 | 0.0000933 | 0.0000918 | 0.0089979 | 0.0093078 | 1.000370 | 2775.502 | 2456.631 |
| K | 78.0779022 | 78.0308500 | 2.0410570 | 2.0913556 | 74.7759800 | 81.4455450 | 1.000474 | 2242.412 | 2490.401 |
| x0[1] | 0.0017213 | 0.0017205 | 0.0000482 | 0.0000495 | 0.0016421 | 0.0018003 | 1.000803 | 1766.489 | 2153.531 |
| x0[2] | 0.0038656 | 0.0038651 | 0.0000825 | 0.0000839 | 0.0037321 | 0.0040025 | 1.001171 | 1639.284 | 2101.449 |
| A0[1] | 78.4312216 | 78.4330500 | 0.1657818 | 0.1696836 | 78.1593950 | 78.6969050 | 1.000606 | 4814.000 | 3311.872 |
| A0[2] | 75.9353760 | 75.9342000 | 0.1969456 | 0.1939982 | 75.6110850 | 76.2584050 | 1.000424 | 4851.169 | 3128.147 |
| sigma_x | 0.0233122 | 0.0233146 | 0.0005440 | 0.0005437 | 0.0224221 | 0.0241935 | 1.001099 | 2614.637 | 2748.456 |
| sigma_A | 0.4006084 | 0.4006225 | 0.0051806 | 0.0052447 | 0.3919497 | 0.4091731 | 1.000296 | 3038.007 | 2443.670 |

**DSMZ-22533**

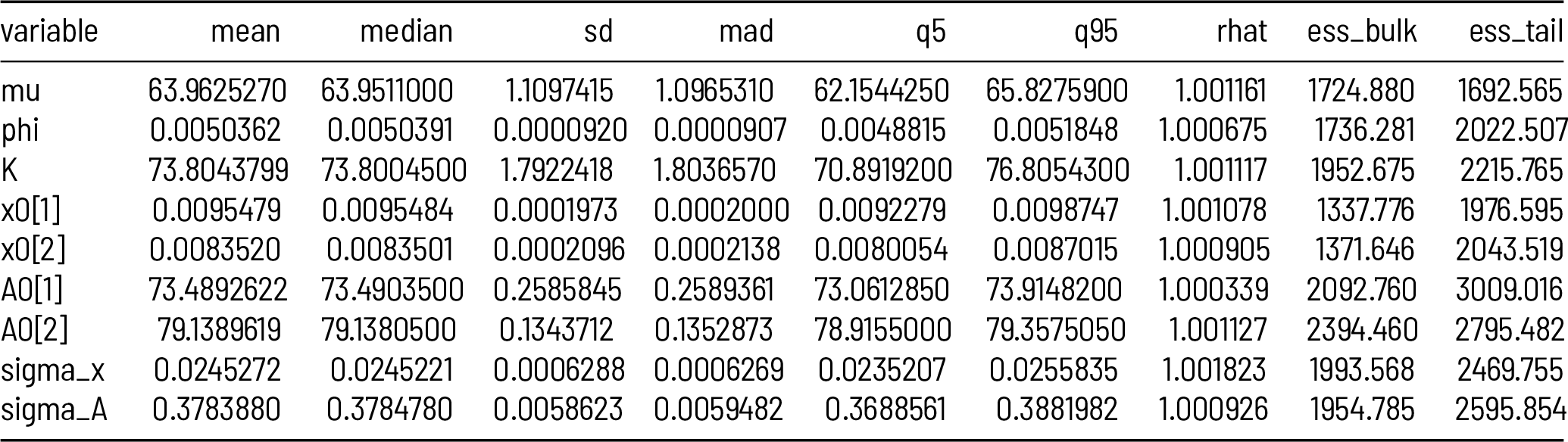

| variable | mean | median | sd | mad | q5 | q95 | rhat | ess_bulk | ess_tail |
| --- | --- | --- | --- | --- | --- | --- | --- | --- | --- |
| mu | 63.9625270 | 63.9511000 | 1.1097415 | 1.0965310 | 62.1544250 | 65.8275900 | 1.001161 | 1724.880 | 1692.565 |
| phi | 0.0050362 | 0.0050391 | 0.0000920 | 0.0000907 | 0.0048815 | 0.0051848 | 1.000675 | 1736.281 | 2022.507 |
| K | 73.8043799 | 73.8004500 | 1.7922418 | 1.8036570 | 70.8919200 | 76.8054300 | 1.001117 | 1952.675 | 2215.765 |
| x0[1] | 0.0095479 | 0.0095484 | 0.0001973 | 0.0002000 | 0.0092279 | 0.0098747 | 1.001078 | 1337.776 | 1976.595 |
| x0[2] | 0.0083520 | 0.0083501 | 0.0002096 | 0.0002138 | 0.0080054 | 0.0087015 | 1.000905 | 1371.646 | 2043.519 |
| A0[1] | 73.4892622 | 73.4903500 | 0.2585845 | 0.2589361 | 73.0612850 | 73.9148200 | 1.000339 | 2092.760 | 3009.016 |
| A0[2] | 79.1389619 | 79.1380500 | 0.1343712 | 0.1352873 | 78.9155000 | 79.3575050 | 1.001127 | 2394.460 | 2795.482 |
| sigma_x | 0.0245272 | 0.0245221 | 0.0006288 | 0.0006269 | 0.0235207 | 0.0255835 | 1.001823 | 1993.568 | 2469.755 |
| sigma_A | 0.3783880 | 0.3784780 | 0.0058623 | 0.0059482 | 0.3688561 | 0.3881982 | 1.000926 | 1954.785 | 2595.854 |

**DSMZ-24814**

| variable | mean | median | sd | mad | q5 | q95 | rhat | ess_bulk | ess_tail |
| --- | --- | --- | --- | --- | --- | --- | --- | --- | --- |
| mu | 42.2652199 | 42.2234000 | 1.1217051 | 1.1162495 | 40.4657700 | 44.1563000 | 1.0002915 | 1640.635 | 1986.899 |
| phi | 0.0067231 | 0.0067269 | 0.0002057 | 0.0002073 | 0.0063786 | 0.0070611 | 1.0010669 | 1579.719 | 2077.161 |
| K | 89.4844345 | 89.4710500 | 2.1338514 | 2.1433948 | 85.9959100 | 93.0136450 | 1.0006271 | 2373.401 | 2341.376 |
| x0[1] | 0.0066060 | 0.0066024 | 0.0001591 | 0.0001569 | 0.0063484 | 0.0068750 | 1.0034752 | 1741.996 | 2147.414 |
| x0[2] | 0.0045008 | 0.0044993 | 0.0001214 | 0.0001187 | 0.0043034 | 0.0047068 | 1.0029807 | 1597.383 | 2270.807 |
| A0[1] | 81.9605948 | 81.9570000 | 0.1898141 | 0.1894021 | 81.6541550 | 82.2738300 | 1.0007350 | 4444.723 | 2901.587 |
| A0[2] | 77.5739450 | 77.5713500 | 0.2705390 | 0.2773945 | 77.1404900 | 78.0154250 | 0.9997288 | 2108.617 | 2515.373 |
| sigma_x | 0.0353289 | 0.0353098 | 0.0006237 | 0.0006220 | 0.0343319 | 0.0363913 | 1.0001487 | 1991.155 | 2369.209 |
| sigma_A | 0.5083501 | 0.5083660 | 0.0058919 | 0.0060601 | 0.4988665 | 0.5180164 | 1.0013341 | 2056.236 | 2498.502 |

**PB-SARUR**

| variable | mean | median | sd | mad | q5 | q95 | rhat | ess_bulk | ess_tail |
| --- | --- | --- | --- | --- | --- | --- | --- | --- | --- |
| mu | 28.1266089 | 28.1179000 | 0.6491801 | 0.6656133 | 27.0818400 | 29.2003100 | 1.0012577 | 1621.421 | 2089.412 |
| phi | 0.0034977 | 0.0034967 | 0.0000931 | 0.0000920 | 0.0033476 | 0.0036540 | 1.0015549 | 1687.955 | 1936.752 |
| K | 0.2500943 | 0.1874790 | 0.2169896 | 0.1884488 | 0.0159445 | 0.6790079 | 1.0018849 | 1683.368 | 1362.333 |
| x0[1] | 0.0001035 | 0.0001036 | 0.0000045 | 0.0000044 | 0.0000961 | 0.0001108 | 1.0014100 | 1735.267 | 1984.098 |
| x0[2] | 0.0034598 | 0.0034587 | 0.0000844 | 0.0000828 | 0.0033222 | 0.0036023 | 1.0011736 | 1644.962 | 2161.432 |
| A0[1] | 80.3749952 | 80.3754000 | 0.1114772 | 0.1114915 | 80.1939900 | 80.5600250 | 1.0007970 | 4096.310 | 2316.685 |
| A0[2] | 79.5229914 | 79.5222500 | 0.1752327 | 0.1764294 | 79.2390650 | 79.8141150 | 1.0000152 | 4101.038 | 2909.671 |
| sigma_x | 0.0219825 | 0.0219814 | 0.0005212 | 0.0005143 | 0.0211322 | 0.0228398 | 0.9997679 | 2674.910 | 2789.782 |
| sigma_A | 0.3412309 | 0.3413015 | 0.0052117 | 0.0052640 | 0.3325613 | 0.3496548 | 1.0009202 | 2891.384 | 2356.799 |

**PB-SCBYV**

| variable | mean | median | sd | mad | q5 | q95 | rhat | ess_bulk | ess_tail |
| --- | --- | --- | --- | --- | --- | --- | --- | --- | --- |
| mu | 32.8331798 | 32.8334000 | 0.5498072 | 0.5679841 | 31.9467750 | 33.7384350 | 1.000523 | 1856.177 | 2171.334 |
| phi | 0.0103116 | 0.0103113 | 0.0001236 | 0.0001239 | 0.0101047 | 0.0105108 | 1.001662 | 2692.683 | 2615.042 |
| K | 88.2649888 | 88.2407000 | 2.1005906 | 2.1308669 | 84.8203000 | 91.6876700 | 1.002180 | 2110.795 | 2311.319 |
| x0[1] | 0.0061185 | 0.0061172 | 0.0001191 | 0.0001213 | 0.0059185 | 0.0063122 | 1.000923 | 1573.290 | 2165.977 |
| x0[2] | 0.0055325 | 0.0055324 | 0.0001283 | 0.0001273 | 0.0053177 | 0.0057382 | 1.000954 | 1629.972 | 2158.675 |
| A0[1] | 80.2987214 | 80.2958000 | 0.2317382 | 0.2297289 | 79.9148450 | 80.6755500 | 1.000589 | 4586.545 | 3065.033 |
| A0[2] | 81.3093830 | 81.3080500 | 0.1464601 | 0.1467774 | 81.0669950 | 81.5484500 | 1.000479 | 6368.718 | 3156.686 |
| sigma_x | 0.0306303 | 0.0306292 | 0.0005185 | 0.0005082 | 0.0297917 | 0.0314976 | 1.000993 | 2936.100 | 2744.984 |
| sigma_A | 0.3974329 | 0.3972985 | 0.0055257 | 0.0056420 | 0.3886542 | 0.4065757 | 1.000837 | 2385.970 | 2659.632 |

**PB-SDVAP**

| variable | mean | median | sd | mad | q5 | q95 | rhat | ess_bulk | ess_tail |
| --- | --- | --- | --- | --- | --- | --- | --- | --- | --- |
| mu | 65.0072534 | 65.0152000 | 0.9424787 | 0.9557581 | 63.4368250 | 66.5292700 | 1.001938 | 1869.036 | 2369.149 |
| phi | 0.0066987 | 0.0066995 | 0.0000779 | 0.0000788 | 0.0065722 | 0.0068269 | 1.002560 | 2803.120 | 2590.320 |
| K | 22.2369412 | 22.2304500 | 0.9774926 | 0.9552392 | 20.6391700 | 23.8884200 | 1.001933 | 2042.293 | 2439.202 |
| x0[1] | 0.0031794 | 0.0031791 | 0.0000722 | 0.0000731 | 0.0030627 | 0.0032990 | 1.002026 | 1533.057 | 2364.353 |
| x0[2] | 0.0038449 | 0.0038465 | 0.0000869 | 0.0000888 | 0.0037015 | 0.0039862 | 1.001551 | 1642.397 | 2461.269 |
| A0[1] | 79.3913281 | 79.3909500 | 0.1089834 | 0.1083039 | 79.2122000 | 79.5715050 | 1.001690 | 5299.491 | 2917.276 |
| A0[2] | 80.8765930 | 80.8777000 | 0.2914804 | 0.2877727 | 80.3876800 | 81.3556200 | 1.000968 | 3092.791 | 2675.625 |
| sigma_x | 0.0161790 | 0.0161640 | 0.0005255 | 0.0005194 | 0.0153238 | 0.0170815 | 1.002377 | 2768.253 | 2529.524 |
| sigma_A | 0.3129388 | 0.3128115 | 0.0050221 | 0.0052232 | 0.3048049 | 0.3211286 | 1.001997 | 3261.594 | 2783.637 |

**PB-SJWFW**

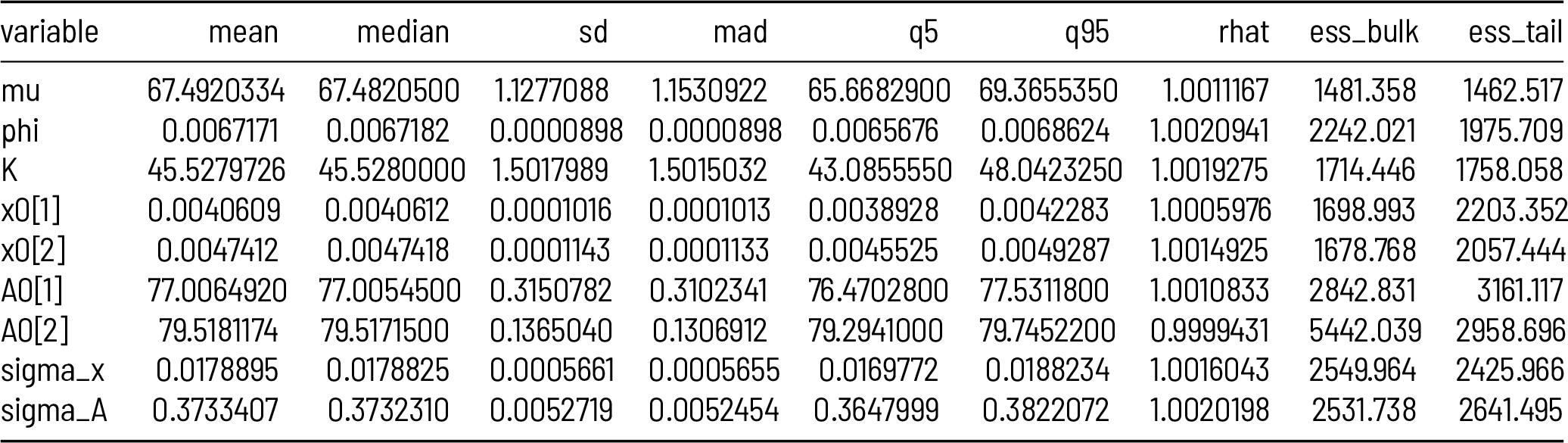

**PB-SPUPY**

| variable | mean | median | sd | mad | q5 | q95 | rhat | ess_bulk | ess_tail |
| --- | --- | --- | --- | --- | --- | --- | --- | --- | --- |
| mu | 59.2841231 | 59.2828500 | 0.8746658 | 0.8848157 | 57.8310850 | 60.7565300 | 1.0016851 | 1646.626 | 1989.842 |
| phi | 0.0063315 | 0.0063307 | 0.0000665 | 0.0000668 | 0.0062234 | 0.0064405 | 1.0015368 | 2646.877 | 2433.222 |
| K | 24.5210095 | 24.5282500 | 1.0301174 | 1.0287020 | 22.8468600 | 26.2351650 | 1.0008388 | 1983.403 | 2389.381 |
| x0[1] | 0.0059946 | 0.0059911 | 0.0001293 | 0.0001296 | 0.0057845 | 0.0062120 | 0.9998119 | 1774.364 | 2348.892 |
| x0[2] | 0.0039506 | 0.0039497 | 0.0000889 | 0.0000893 | 0.0038078 | 0.0041019 | 1.0002608 | 1610.042 | 2343.163 |
| A0[1] | 77.5003594 | 77.5009500 | 0.2023349 | 0.2022266 | 77.1663700 | 77.8340450 | 1.0020316 | 4478.333 | 2658.752 |
| A0[2] | 77.1270053 | 77.1277000 | 0.0951279 | 0.0945899 | 76.9711850 | 77.2816150 | 1.0007736 | 4264.951 | 3100.926 |
| sigma_x | 0.0129983 | 0.0130046 | 0.0005369 | 0.0005327 | 0.0121290 | 0.0139122 | 1.0008532 | 2209.255 | 2164.781 |
| sigma_A | 0.2037247 | 0.2037395 | 0.0051985 | 0.0052017 | 0.1950968 | 0.2121761 | 1.0021472 | 2950.317 | 2579.972 |

**PB-SSJQB**

| variable | mean | median | sd | mad | q5 | q95 | rhat | ess_bulk | ess_tail |
| --- | --- | --- | --- | --- | --- | --- | --- | --- | --- |
| mu | 38.0671042 | 38.0624500 | 0.8699528 | 0.8638369 | 36.6803750 | 39.5398100 | 1.002388 | 1656.544 | 1825.423 |
| phi | 0.0078834 | 0.0078894 | 0.0001885 | 0.0001860 | 0.0075681 | 0.0081869 | 1.002370 | 1803.847 | 2029.734 |
| K | 88.6392096 | 88.6650000 | 2.1131475 | 2.0969153 | 85.1171350 | 92.0992250 | 1.001670 | 2048.691 | 2447.895 |
| x0[1] | 0.0040742 | 0.0040716 | 0.0001054 | 0.0001057 | 0.0039032 | 0.0042474 | 1.003933 | 1448.427 | 2014.783 |
| x0[2] | 0.0070930 | 0.0070900 | 0.0001571 | 0.0001605 | 0.0068398 | 0.0073515 | 1.003282 | 1579.399 | 2003.844 |
| A0[1] | 75.9330331 | 75.9323500 | 0.2427746 | 0.2472977 | 75.5380300 | 76.3378250 | 1.002558 | 2318.624 | 2662.199 |
| A0[2] | 81.2745600 | 81.2780000 | 0.1775958 | 0.1815444 | 80.9800950 | 81.5627100 | 1.001505 | 4674.038 | 2987.650 |
| sigma_x | 0.0341382 | 0.0341277 | 0.0006036 | 0.0006121 | 0.0331840 | 0.0351560 | 1.001816 | 2003.229 | 1899.752 |
| sigma_A | 0.4694522 | 0.4693675 | 0.0058128 | 0.0057592 | 0.4598969 | 0.4790814 | 1.002598 | 2250.525 | 2523.170 |

**Model 2**

**DSMZ-24814**

Model 2 DSMZ-24814
| variable | mean | median | sd | mad | q5 | q95 | rhat | ess_bulk | ess_tail |
| --- | --- | --- | --- | --- | --- | --- | --- | --- | --- |
| mu | 12.4578546 | 12.4583500 | 0.2947238 | 0.2911826 | 11.9734900 | 12.9461450 | 1.0012667 | 2555.423 | 2622.631 |
| phi | 0.0014204 | 0.0014167 | 0.0001311 | 0.0001350 | 0.0012106 | 0.0016377 | 1.0013102 | 2237.722 | 2598.371 |
| K | 55.4279843 | 55.3966000 | 2.0013997 | 1.9983224 | 52.1136500 | 58.7198450 | 1.0002784 | 2689.050 | 2528.263 |
| mu2 | 91.3383973 | 91.2727500 | 2.2224156 | 2.1999560 | 87.7950950 | 95.1448400 | 1.0022282 | 2266.863 | 2434.473 |
| phi2 | 0.0044903 | 0.0044931 | 0.0000769 | 0.0000765 | 0.0043603 | 0.0046138 | 1.0014151 | 2115.389 | 2714.317 |
| K2 | 35.6677337 | 35.6422500 | 2.3639028 | 2.3363552 | 31.7553900 | 39.6394500 | 1.0003332 | 3265.636 | 2675.488 |
| x0[1] | 0.0036464 | 0.0036474 | 0.0000890 | 0.0000876 | 0.0034987 | 0.0037932 | 0.9995410 | 3121.885 | 2633.438 |
| x0[2] | 0.0020883 | 0.0020899 | 0.0000714 | 0.0000726 | 0.0019697 | 0.0022041 | 1.0002947 | 2754.908 | 2257.442 |
| A0[1] | 80.6813264 | 80.6823500 | 0.1194130 | 0.1201647 | 80.4832950 | 80.8785150 | 0.9998932 | 5821.417 | 3395.289 |
| A0[2] | 79.8959982 | 79.8994500 | 0.2006775 | 0.2041540 | 79.5685900 | 80.2218150 | 1.0006110 | 4076.876 | 3475.659 |
| sigma_x | 0.0178874 | 0.0178874 | 0.0005053 | 0.0004884 | 0.0170505 | 0.0187348 | 1.0010506 | 3849.580 | 2561.786 |
| sigma_A | 0.3275759 | 0.3275460 | 0.0051208 | 0.0051417 | 0.3191967 | 0.3361212 | 1.0010588 | 4405.287 | 2388.663 |

**PB-SCBYV**

| variable | mean | median | sd | mad | q5 | q95 | rhat | ess_bulk | ess_tail |
| --- | --- | --- | --- | --- | --- | --- | --- | --- | --- |
| mu | 11.1899534 | 11.1796000 | 0.2938624 | 0.2958528 | 10.7160800 | 11.6945350 | 1.0016780 | 2075.715 | 2257.250 |
| phi | 0.0059236 | 0.0059267 | 0.0001757 | 0.0001781 | 0.0056300 | 0.0062054 | 1.0011842 | 1904.726 | 2449.120 |
| K | 41.2279974 | 41.1751500 | 1.7655031 | 1.7522849 | 38.3304900 | 44.1698150 | 1.0014193 | 2199.143 | 2457.095 |
| mu2 | 106.3820994 | 106.3505000 | 3.6302076 | 3.6027180 | 100.4576500 | 112.3920500 | 1.0012945 | 2014.978 | 2444.193 |
| phi2 | 0.0035055 | 0.0035028 | 0.0001066 | 0.0001058 | 0.0033365 | 0.0036821 | 1.0013390 | 1796.219 | 2321.114 |
| K2 | 31.9677349 | 31.9601000 | 2.6348501 | 2.6219040 | 27.6349100 | 36.3320150 | 1.0008010 | 3352.031 | 2546.642 |
| x0[1] | 0.0039724 | 0.0039717 | 0.0000899 | 0.0000909 | 0.0038249 | 0.0041223 | 1.0003016 | 3161.437 | 3043.678 |
| x0[2] | 0.0047753 | 0.0047762 | 0.0000972 | 0.0000959 | 0.0046153 | 0.0049332 | 1.0010805 | 3401.187 | 3143.707 |
| A0[1] | 79.7747086 | 79.7742000 | 0.2275675 | 0.2284687 | 79.4028200 | 80.1570050 | 1.0019512 | 3521.253 | 3244.725 |
| A0[2] | 80.5357060 | 80.5357500 | 0.1164228 | 0.1160876 | 80.3425900 | 80.7260150 | 0.9999303 | 5246.264 | 3553.522 |
| sigma_x | 0.0194494 | 0.0194428 | 0.0004828 | 0.0004981 | 0.0186759 | 0.0202615 | 1.0004661 | 3870.444 | 2735.458 |
| sigma_A | 0.3121337 | 0.3121110 | 0.0051310 | 0.0050779 | 0.3036693 | 0.3205572 | 1.0012656 | 3990.732 | 2466.391 |

**PB-SSJQB**

| variable | mean | median | sd | mad | q5 | q95 | rhat | ess_bulk | ess_tail |
| --- | --- | --- | --- | --- | --- | --- | --- | --- | --- |
| mu | 10.8089404 | 10.8037000 | 0.2793707 | 0.2755412 | 10.3547550 | 11.2835050 | 1.0004861 | 1957.430 | 2476.664 |
| phi | 0.0039730 | 0.0039740 | 0.0001540 | 0.0001516 | 0.0037196 | 0.0042309 | 0.9999424 | 1914.839 | 1967.771 |
| K | 41.5173065 | 41.4756500 | 1.8119793 | 1.8097357 | 38.6404650 | 44.5094000 | 1.0006654 | 2092.014 | 2457.911 |
| mu2 | 133.8975512 | 133.8380000 | 4.0850327 | 4.0400850 | 127.1798500 | 140.8407500 | 1.0002956 | 2187.607 | 2636.377 |
| phi2 | 0.0034129 | 0.0034140 | 0.0000856 | 0.0000850 | 0.0032674 | 0.0035530 | 1.0006044 | 1874.257 | 1976.615 |
| K2 | 33.5031737 | 33.4698000 | 2.5114189 | 2.5479964 | 29.3962750 | 37.7073700 | 1.0019368 | 2961.051 | 2396.859 |
| x0[1] | 0.0014556 | 0.0014548 | 0.0000629 | 0.0000618 | 0.0013541 | 0.0015611 | 1.0001717 | 2857.669 | 2205.682 |
| x0[2] | 0.0042749 | 0.0042731 | 0.0001036 | 0.0001028 | 0.0041074 | 0.0044459 | 1.0007601 | 3329.335 | 2791.551 |
| A0[1] | 80.2111927 | 80.2074000 | 0.2333630 | 0.2317304 | 79.8297850 | 80.5999450 | 1.0001299 | 3926.406 | 2850.345 |
| A0[2] | 80.1582466 | 80.1576500 | 0.1232057 | 0.1223886 | 79.9564000 | 80.3639050 | 1.0004059 | 4281.315 | 3464.883 |
| sigma_x | 0.0189369 | 0.0189256 | 0.0005429 | 0.0005224 | 0.0180630 | 0.0198435 | 1.0003841 | 3833.573 | 2600.213 |
| sigma_A | 0.3139960 | 0.3138770 | 0.0052837 | 0.0051913 | 0.3054359 | 0.3228022 | 1.0007950 | 3179.629 | 2547.788 |

